# Aurora kinase A is a synthetic lethal target in FANCA-deficient cancers

**DOI:** 10.64898/2026.02.04.703705

**Authors:** Dilara Akhoundova, Martín González-Fernández, Aavash Baral, Andrej Benjak, Carmen Perry, Dinda Shezaria Hardy Lubis, Ladina Hörtensteiner, Saskia Hussung, Lea Lingg, Sina Maletti, Aino Paasinen-Sohns, Marika Lehner, Bastian Dislich, Erik Vassella, Simone de Brot, Therese Waldburger, Petros Tsiridis, Carmen Cardozo, Ralph Fritsch, Clelia Pistoni, Seynabou Diop, Phillip Thienger, Taina Kaiponen, Muriel Jaquet, Mariana Ricca, Smruthy Sivakumar, Christopher McDermott-Roe, David E. Root, Paola Francica, Sven Rottenberg, Mark A. Rubin

**Affiliations:** Department of Medical Oncology, Inselspital, University Hospital of Bern, University of Bern, Bern, 3010 Switzerland; Department for BioMedical Research, University of Bern, Bern, 3008 Switzerland; Bern Center for Precision Medicine, Inselspital, University Hospital of Bern, Bern, 3008 Switzerland; Institute of Animal Pathology, Vetsuisse Faculty, University of Bern, 3012 Bern, Switzerland; Institute of Tissue Medicine and Pathology, University of Bern, Bern, 3008 Switzerland; Department of Medical Oncology and Hematology, University Hospital of Zurich, University of Zurich, Zurich, 8091 Switzerland; Cancer Genomics Research, Foundation Medicine Inc., Boston, Massachusetts, US; Genetic Perturbation Platform, Broad Institute, Cambridge, Massachusetts, US

**Author notes:** Co-corresponding authors: Prof Mark A. Rubin, M.D. Murtenstrasse 24, 3008 Bern, Switzerland, Dilara Akhoundova, M.D. Freiburgstrasse 41, Lory-Haus, 3008 Bern, Switzerland. equal contribution (Dilara Akhoundova, Martín González-Fernández). Declaration of interests: S.S. is an employee at Foundation Medicine, Inc., with equity interest in Roche.

**Keywords:** Aurora kinase A (AURKA), FANCA, Fanconi anemia pathway, synthetic lethality, mitosis, mitotic spindle, cell cycle, prostate cancer, head-and-neck cancer, precision oncology

## Abstract

Loss-of-function genomic alterations in *FANCA* occur across multiple cancer types, yet no molecularly tailored therapies have successfully exploited this potential vulnerability. Using complementary unbiased approaches, including a genome-wide CRISPR/Cas9 loss-of-function screen and a high-throughput drug screen in isogenic cancer cell-based models, we identified Aurora kinase A (AURKA) as a reproducible synthetic lethal target of FANCA-deficient cancers. Inhibition of AURKA induced chromosomal instability, micronucleation, and early G2/M arrest selectively in FANCA-deficient cells, consistent with an increased reliance on mitotic checkpoint control. Mechanistically, FANCA deficiency is associated with an elevated AURKA expression at both the transcriptomic and protein levels, and with an upregulation of mitotic spindle and G2/M checkpoint gene signatures. Analysis of large-scale cancer genomics datasets, including over 650,000 clinically sequenced tumors, confirms that *FANCA* is the most frequently altered Fanconi anemia pathway gene across cancers, and that Fanconi anemia–defective tumors exhibit an increased tumor mutational burden and genomic instability. Collectively, our findings point to AURKA inhibition as a promising precision treatment strategy in FANCA-deficient cancers and provide a rationale to further explore this strategy in the clinic.

## 1. Introduction

Genomic alterations in *FANCA* and other core components of the Fanconi Anemia (FA) pathway occur across multiple cancer types, affecting a substantial number of patients. Despite a large body of mechanistic work dissecting the pathogenetic role of the FA pathway in tumor initiation and progression, no successful molecularly targeted treatment strategies exist for FA pathway-altered malignancies. In the clinical scenario, *FANCA* and other FA pathway gene alterations have been most extensively studied in prostate cancer. Here, several trials evaluating poly (ADP-ribose)-polymerase (PARP) inhibitors, including the recently published AMPLITUDE trial, failed to show a consistent benefit in FA-altered tumors, emphasizing the need for alternative molecularly targeted treatment strategies ^1–5^.

The classical role of the FA complex is to repair toxic interstrand crosslinks (ICLs), which lead to stalling of the DNA replication fork and double-strand breaks (DSBs) during the S phase of the cell cycle ^6,7^. ICLs are detected by the FANCM–FAAP24–MHF1–MHF2 complex, which subsequently recruits further proteins of the FA core complex, including FANCA, to ICL sites ^7^. Activation of the FA core complex enables the monoubiquitination of FANCD2 and FANCI heterodimers, promoting nucleotide excision by the ERCC4-ERCC1 complex, required to unhook the ICL from one of the parental DNA strands ^7^. The crosslinked nucleotide, tethered to the complementary strand, is then bypassed by error-prone translesion synthesis polymerases, such as REV1 or DNA polymerase ζ (REV3–REV7). Once translesion synthesis is completed, the intact DNA duplex serves as a template for homologous recombination-mediated repair (HRR) of the DSB induced by the incisions. The BRCA2-PALB2 complex, together with RAD51, plays an essential role in the HRR pathway, which concludes the FA pathway-mediated ICL repair^8^. Beyond ICL repair, FANCA is also involved in other DNA damage repair (DDR) pathways, such as the single-strand annealing pathway ^7,9,10^.

In addition to the well-characterized role in ICL repair, recent research points to a role of FA pathway members in mitotic spindle regulation and DNA repair during mitosis, found to be especially relevant at centromeric regions ^11–13^. Moreover, FANCA has been shown to regulate mitotic spindle assembly during early mitosis and to interact with centromeric protein CENP-E. Loss of FANCA leads to lagging chromosomes and micronuclei formation; however, vulnerability to antimitotic drugs is underexplored ^11,14,15^. Lastly, the intact FA complex has been shown to drive chromothripsis of mis-segregated chromosomes from micronuclei during mitosis ^16^.

Aurora kinase A (AURKA) is a key mitotic kinase, regulating mitotic spindle assembly, centrosome maturation, and mitotic entry ^17,18^. AURKA expression is cell cycle-dependent, peaking during G2/M and undergoing a complex multi-level regulation on the transcriptional, post-transcriptional, translational, and post-translational levels ^19^. During mitotic entry, AURKA cooperates with other mitotic kinases such as Polo-like kinase 1 (PLK1) ^20,21^. PLK1 orchestrates mitotic timing downstream of AURKA by regulating CDK1, APC/C, and Cyclin B1, further contributing to bipolar spindle assembly and correct kinetochore-microtubule attachments. In cancer, *AURKA* amplification and/or overexpression has been widely reported to promote chromosomal instability (CIN), an aggressive tumor phenotype, and therapy resistance ^22^. Previous reports have shown some links between members of the FA pathway and this mitotic kinase axis: AURKA is required for the activation of the FA pathway in response to DNA damage ^23^, and FA-deficient cell models are more sensitive to PLK1 inhibitors^24^. Degradation of FANCM in mitosis is also orchestrated by PLK1 ^12^. Moreover, analysis of gene expression signatures from FANCA-deficient vs -proficient squamous cell head-and-neck cancers showed upregulation of the centriole duplication signature ^25^. Together, these observations suggest that disruption of the FA pathway may increase cellular reliance on mitotic kinase signaling to preserve chromosome segregation fidelity, raising the possibility that FANCA-deficient cancers acquire targetable dependencies within the AURKA–PLK1 mitotic axis.

In this study, we sought to systematically identify actionable vulnerabilities arising from FANCA deficiency using complementary unbiased functional approaches. By combining a genome-wide CRISPR/Cas9 loss-of-function synthetic lethality screen with a high-throughput small-molecule drug screen in isogenic FANCA-proficient and -deficient cancer models, we uncovered a reproducible dependency on AURKA. We show that FANCA loss is associated with elevated AURKA expression, increased mitotic stress, and heightened reliance on G2/M checkpoint and mitotic spindle regulation. Pharmacological inhibition of AURKA selectively compromises the viability of FANCA-deficient cells by exacerbating CIN and disrupting mitotic progression. Together with analyses of large-scale cancer genomics and transcriptomic datasets, our findings define a mitosis-centred synthetic lethal vulnerability in FANCA-deficient cancers and provide a biologically informed framework for therapeutic targeting of this molecular subset.

## Results

### Genome-wide loss-of-function CRISPR/Cas9 and high-throughput drug screens nominate AURKA as a synthetically lethal target in FANCA-deficient cancers

To identify genes whose loss is synthetically lethal with FANCA-deficiency, we performed a genome-wide loss-of-function screen in previously generated isogenic human androgen-independent prostate cancer FANCA-deficient vs -proficient DU145 cell lines, using the Brunello all-in-one Hygromycin whole-genome library (Fig. 1A). A detailed description of the screen design is provided in the methods section. Briefly, transduced cells were seeded and then cultured for 6 days. Genomic DNA was then extracted and sgRNA abundance was measured via next-generation sequencing (NGS). We performed the screen in biological duplicates to increase robustness. Quality control parameters indicated good replicate correlation and consistent depletion of essential genes (Suppl. Fig. 1A-C). MAGeCK and MAGeCKFlute were used to identify differentially represented sgRNAs and biological processes, respectively. For the purpose of this manuscript, we focused on genes whose loss reduced viability in the context of FANCA deficiency. Our screen identified multiple genes coding for proteins involved in mitosis. Depleted hits included genes regulating mitotic spindle assembly (*AURKA, HAUS1, HAUS2, HAUS6, HAUS7, DYNC1H1, KIF18B*), DNA repair during mitosis via microhomology-mediated end-joining (MMEJ) (*911 complex: RAD1*, *HUS1, RAD17*), DNA damage checkpoint (*CLSPN, CDC6, CHEK1, ATRIP, RNF8, KAT5, CCNB1*), chromosome condensation (*SMC2, SMC4, NCAPG, KMT5A*) and kinetochore regulation (*NDC80, TTK, BUB1, BUB1B, INCENP, CENPC, CENPT)* (Fig. 1B). In line with these findings, pathway enrichment analysis showed depletion in targets involved in cell division and G2/M checkpoint (Fig. 1C, D, Suppl. Fig. 2A,B). As a complementary unbiased approach, isogenic FANCA-deficient vs -proficient DU145 cell lines were exposed to a semi-automatic high-throughput drug screen using a DNA repair and cell cycle dedicated library (1624 compounds) (Fig. 1E). In line with the CRISPR screen, top drugs leading to differential sensitivity (effect size >0.3 and statistically significant difference) included drugs targeting AURKA (CD532), PLK1 (BI-2536) or other targets involved in mitosis (IC261). The latter has been shown to inhibit microtubule polymerization, probably in a Casein kinase 1 delta and epsilon (CK1δ/ɛ)-independent manner ^26,27^ (Fig. 1F). To validate our results in a real FA-patient derived head-and-neck squamous carcinoma cell line, we used the CCH-SCC-FA1 (*FANCA*^⁻/⁻^) cell line, in comparison with the isogenic FANCA complemented line CCH-SCC-FA1+Tg(LV) (*FANCA^Compl^*), hereafter referred as CCH-SCC-FA1 (*FANCA^Compl^),* as well as further isogenic *FANCA* -deficient vs -proficient cell pairs. Extensive characterization of these isogenic cell pairs confirmed FANCA protein loss (Suppl. Fig. 3A), lack of FANCD2 monoubiquitination under treatment with the ICL-inducing agents mitomycin C (MMC) and cisplatin (Suppl. Fig. 3B, C), as well as increased sensitivity to treatment with MMC and cisplatin (Suppl. Fig. 3D-G). Impaired FANCD2 foci formation following exposure to MMC was observed in FANCA-deficient cell models (Suppl. Fig. 4A-F). Together, these two orthogonal unbiased screening approaches converged on mitotic regulation as a vulnerability of FANCA-deficient cancer cells, with AURKA being a top-ranked and pharmacologically actionable synthetic lethal candidate.

**Figure 1.**
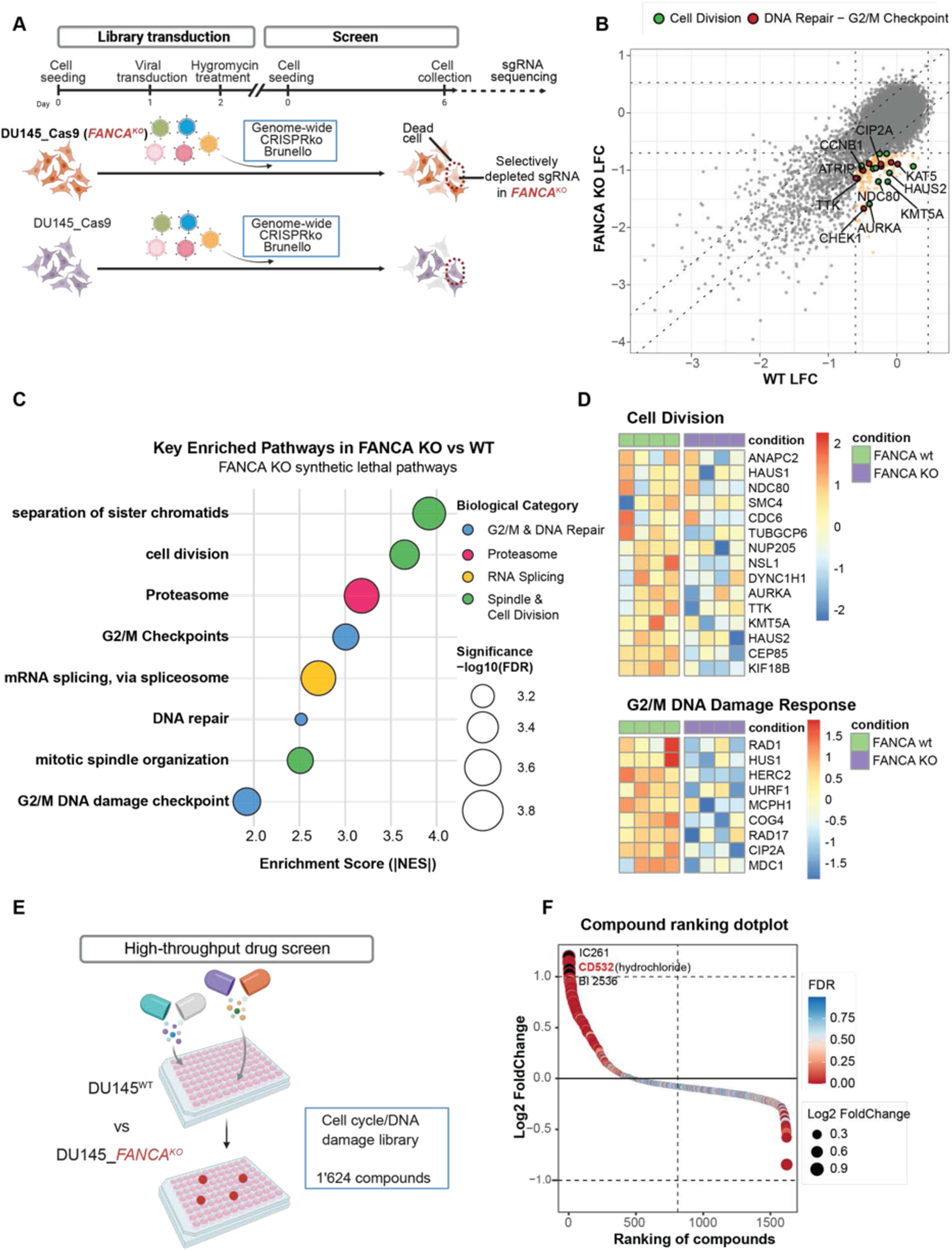
Design and results from a synthetic lethality loss-of-function whole-genome CRISPR/Cas9 and semi-automated high-throughput drug screen performed on isogenic DU145^WT^ and DU145_*FANCA*^KO^ cell lines. (A) Design and read-out of the synthetic lethality loss-of-function whole-genome CRISPR/Cas9 screen performed on isogenic FANCA-proficient vs -deficient DU145 cells. (B) Nine square plot highlighting *AURKA* and other selected differentially depleted hits from the loss-of-function CRISPR screen performed on DU145^WT^ and DU145_*FANCA*^KO^ cell lines, hits were identified through the MAGeCK algorithm. (C) Functional enrichment analysis from the CRISPR screen indicates depletion of targets involved in cell division and G2/M checkpoint. (D) Heatmaps illustrating normalized log fold change (LFC) to Day 0 of sgRNAs targeting cell division and G2/M DNA damage response-related genes from the CRISPR/Cas9 whole genome synthetic lethality screen performed on isogenic FANCA-proficient vs -deficient DU145 cells. (E) Design and schematic illustration of the high-throughput drug screen performed, as a complementary approach, on the same FANCA-proficient vs -deficient DU145 cell lines, a cell cycle and DNA damage-dedicated drug library was used. (F) Results from the high-throughput drug screen show overlapping hits, identifying the AURKA inhibitor CD532, the PLK1 inhibitor BI 2536, and IC261, a microtubule polymerization inhibitor, as top differential hits. *Abbreviations: FDR: false discovery rate; NES: normalized enrichment score.* Figure 1A *was created using BioRender*.

### Inhibition of AURKA is synthetically lethal with FANCA-deficiency *in vitro*

Clonogenic growth assays showed increased vulnerability of the FANCA-deficient FA-patient-derived head-and-neck squamous carcinoma CCH-SCC-FA1 (*FANCA^-/-^*) cells, as compared to the isogenic FANCA-complemented CCH-SCC-FA1 (*FANCA^Compl^),* to the AURKA inhibitor alisertib (Fig. 2A,B), as well as to CD532, a second AURKA inhibitor (Fig. 2C,D). Drug response assays confirmed increased sensitivity to alisertib of the CCH-SCC-FA1 (*FANCA*^⁻/⁻^) cells (Fig. 2E). Increased sensitivity to alisertib was also observed in the UM-SCC-01(*FANCA^KO^*) head-and-neck squamous carcinoma cells, as compared to the wild-type (WT) UM-SCC-01 (*FANCA*^+/+^) (Suppl. Fig. 5A). To assess whether FANCA-deficient cells exhibit increased AURKA expression, we assessed AURKA protein levels by Western blotting and immunohistochemistry (IHC), observing increased expression in FANCA-deficient cells (Fig. 2F-H). Increased IHC levels were confirmed on the RPE1 hTERT *p53*^- /-^_*FANCA*^KD^ cell line as compared to the corresponding WT line, used as a complementary isogenic cell model (Suppl. Fig. 5B). Following this finding, we assessed mRNA levels by real-time quantitative polymerase chain reaction (RT-qPCR), observing increased basal AURKA gene expression levels in unsynchronized FANCA-deficient CCH-SCC-FA1 (*FANCA*^⁻/⁻^) cells (Suppl. Fig. 5C). As both CRISPR and drug screens pointed to additional mitotic targets, we additionally assessed differential response to other drugs: PLK1 inhibitors (volasertib, BI2536), IC261, and docetaxel. As expected, CCH-SCC-FA1 (*FANCA*^⁻/⁻^) cells were hypersensitive to all these drugs (Suppl. Fig. 6A-J). These data demonstrate that FANCA loss confers a selective and reproducible hypersensitivity to AURKA inhibition across several isogenic models, accompanied by increased AURKA expression and a broader vulnerability to perturbations of mitotic control.

**Figure 2.**
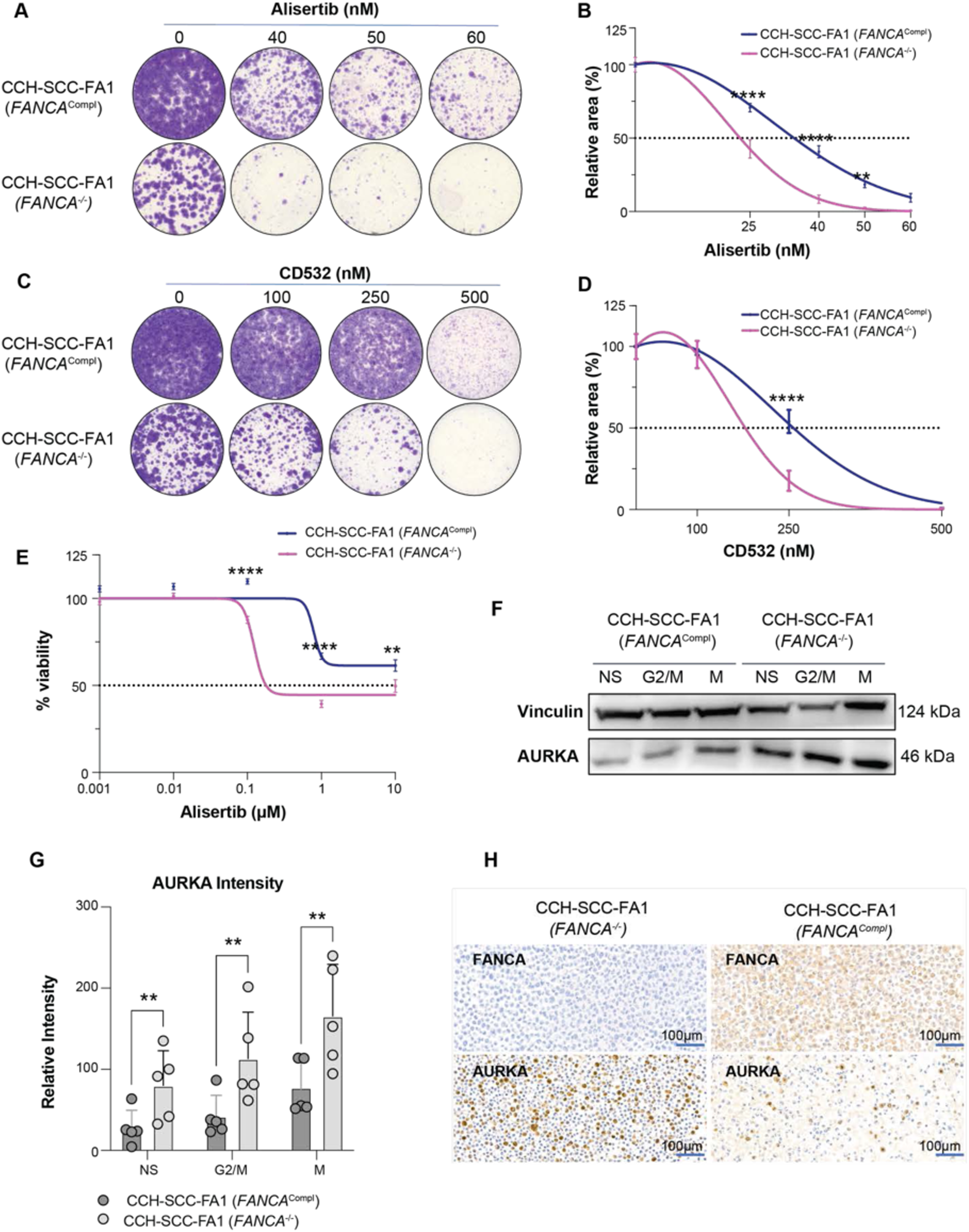
***In vitro* hit validation from CRISPR/Cas9 whole-genome synthetic lethality and high-throughput drug screens confirms that FANCA loss is synthetically lethal with AURKA inhibition.** (A-D) Clonogenic growth assays for isogenic squamous head-and-neck FANCA-deficient vs -proficient cancer cell lines CCH-SCC-FA1 (*FANCA*^-/-^) and CCH-SCC-FA1 (*FANCA*^Compl^) under treatment with two AURKA inhibitors, alisertib (A,B) and CD532 (C,D): representative images (A,C) and relative area (B,D) are represented. Statistical significance was calculated with 2-way ANOVA followed by Šidák’s multiple comparison test. (E) Drug response curves, assessed by CellTiter-Glo luminescence cell viability assay, illustrating increased vulnerability to the AURKA inhibitor alisertib of CCH-SCC-FA1 (*FANCA*^-/-^), as compared to CCH-SCC-FA1 (*FANCA*^Compl^) cells. Statistical significance was calculated using the 2-way ANOVA followed by Bonferroni’s multiple comparison test. (F) Representative Western blot showing increased AURKA protein expression in CCH-SCC-FA1 (*FANCA*^- /-^) vs CCH_SCC_FA1 (*FANCA*^Compl^) cells, in non-synchronized (NS) cells, as well as in cells synchronized in G2/M and early mitosis (M). (G) Quantification of AURKA protein expression normalized to vinculin in CCH-SCC-FA1 (*FANCA*^-/-^) vs CCH_SCC_FA1 (*FANCA*^Compl^) cells from 5 biological Western blot replicates in non-synchronized (NS) cells, as well as in cells synchronized in G2/M and early mitosis (M). Mean and standard deviation are depicted, and statistical analysis was performed using a two-way ANOVA and Tuckey’s post-hoc test. **p < 0.02. (H) Increased AURKA protein IHC expression shown on CCH-SCC-FA1 (*FANCA*^-/-^) vs CCH-SCC-FA1 (*FANCA*^Compl^) cell lines, scale bar represents 100 µm.

### Treatment with alisertib enhances CIN and leads to early G2/M arrest in FANCA-deficient cell models

Upon alisertib treatment, we observed increased micronuclei in CCH-SCC-FA1 (*FANCA*^⁻/⁻^) vs CCH-SCC-FA1 (*FANCA^Compl^*) cells (Fig. 3A, B), an indicator of CIN. To distinguish between structural or numerical CIN, we additionally quantified the presence of CENP-B foci in micronuclei, a centromere-associated protein. As shown in Suppl. Fig. 7A, we observed a slight increase in the ratio of CENP-B positive micronuclei between the two cell lines in basal conditions, but only a trend under exposure to alisertib. Interestingly, both cell lines exhibited extensive multinucleation under treatment with alisertib, which was more pronounced in FANCA-proficient cells (Suppl. Fig. 7B, C). Alisertib has been shown to induce G2/M arrest by preventing mitotic entry ^28^, therefore, we sought to assess differences in cell cycle distribution under treatment with alisertib. Alisertib induced G2/M arrest in FANCA-deficient cells earlier than in their FANCA-proficient counterparts. In contrast, a larger fraction of FANCA-proficient cells remained in G1 at 24 and 36 hours (Fig. 3C,D, Suppl. Fig. 7D). When analyzing bulk RNA sequencing data, we observed an upregulation of the G2/M checkpoint hallmark in CCH-SCC-FA1 (*FANCA*^⁻/⁻^) as compared to CCH-SCC-FA1 (*FANCA^Compl^*) cells on the transcriptional level in basal conditions (Suppl. Fig. 8A) as well as under exposure to alisertib (Fig. 3E). We also found higher basal AURKA expression in FANCA-deficient cells, which increased under treatment with alisertib correlating with the above mentioned enhanced G2/M accumulation (Fig. 3F). Treatment with alisertib also upregulated additional centrosome-involved genes in FANCA-deficient cells (*CEP131*, *KIF15*, *TOP2A*) (Suppl. Fig. 8B). These findings indicate that AURKA inhibition exacerbates pre-existing mitotic stress in FANCA-deficient cells, leading to enhanced CIN and an early G2/M arrest that reflects increased reliance on mitotic checkpoint control.

**Figure 3.**
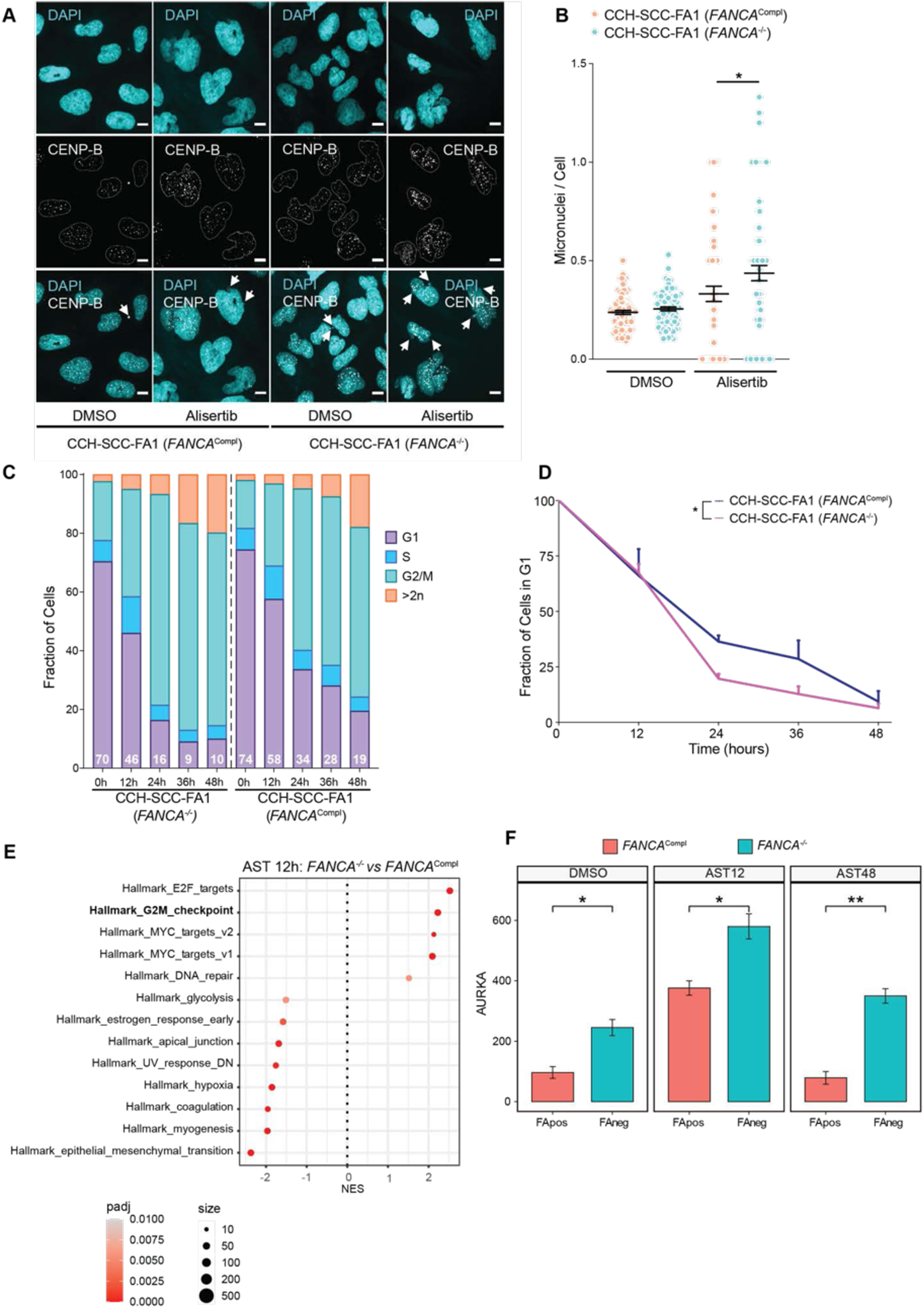
Treatment with alisertib leads to increased mitotic alterations in the presence of FANCA loss. (A,B) Increased micronuclei formation upon treatment with alisertib (500 nM) in CCH-SCC-FA1 (*FANCA*^-/-^) vs CCH-SCC-FA1 (*FANCA*^Compl^) cells: representative immunofluorescence images showing the centromeric marker CENP-B and DAPI. Scale represents 10 µm. (A) and micronuclei quantification in non-multinucleated cells from at least 3 biological replicates. Statistical significance was assessed using a one-way ANOVA followed by Šidák’s multiple comparison test (B). (C) Cell cycle analysis under treatment with alisertib 500 nM over time (0, 12, 24, 36, and 48 hours) in FANCA-deficient vs -proficient CCH-SCC-FA1 cells, quantification from 3 biological replicates. (D) Normalized fraction of FANCA-deficient vs proficient CCH-SCC-FA1 cells remaining in G1 at 0, 12, 24, 36, and 48 hours of alisertib treatment; for statistical significance, the paired t-test was used. (E) Functional enrichment analysis of bulk RNA sequencing from CCH-SCC-FA1 (*FANCA*^-/-^) vs CCH-SCC-FA1 (*FANCA*^Compl^) after 12-hour treatment with alisertib reveals increased expression of the G2/M hallmark in FANCA-deficient cells. (F) Upregulation of AURKA expression under alisertib treatment: higher basal, 12 h, and 48 h levels are observed in CCH_SCC_FA1 (*FANCA*^-/-^) vs CCH_SCC_FA1 (*FANCA*^Compl^) cells. *Abbreviations: FA neg.: CCH-SCC-FA1 (FANCA^-/-^), FA pos.: CCH-SCC-FA1 (FANCA*^Compl^*), NES: normalized enrichment score; padj: adjusted p value*.

### Prevalence and molecular characteristics of FA pathway genomic alterations in human cancer

To assess the potential clinical relevance and prevalence of FANCA-associated tumor vulnerabilities, we next analyzed the genomic landscape and molecular characteristics of Fanconi anemia pathway alterations across a large pan-cancer cohort. For this purpose, we used a pan-cancer dataset of 669,135 cancers molecularly profiled with Foundation Medicine (FMI) targeted NGS assays: FoundationOne^®^, FoundationOne^®^CDx and laboratory developed test FoundationOne^®^Heme, between December 2013 and September 2025, including solid and hematological cancers. After exclusion of microsatellite instable (MSI-high) samples, a cohort of 654,634 tumors remained for analyses. Most frequent histologic tumor subtypes included in the cohort were non-small cell lung cancer (NSCLC) (19%), colorectal (11%), breast (9%), pancreatic (6%), prostate (5%) and ovarian (4%) cancers (Fig. 4A). Globally, FA pathway gene alterations were identified in 11,709 (~1.8%) patients, most frequently affecting *FANCA* (n=6,018, 51% of the cases), followed by *FANCC* (n=2,828, 24%), *FANCG* (n=1,559, 13%) and *FANCL* (n=1,405, 12%), with some cases also exhibiting several of these gene alterations (Fig. 4A, Suppl. Table 1). Genomic alterations identified in *FANCA* were most frequently single nucleotide variants (68%), followed by homozygous deletions (17%) and rearrangements (15%) (Fig. 4A, Suppl. Table 2). A larger proportion of tumor samples with FA gene alterations had high tumor mutational burden (TMB): 24 vs 14% (p<0.001) (Fig. 4B), as well as homologous recombination deficiency (HRD) signature: 12 vs 7 % (p<0.001) (Suppl. Fig. 9A), which is in line with the observed higher CIN in FANCA-deficient cancers (Fig. 3B). Pathogenic or likely pathogenic alterations in *FANCA* were most commonly identified in skin (2.0%), bladder (1.5%), cervix (1.3%), liver (1.1%), salivary gland (1.1%), breast (1.1%), NSCLC (1.1%), head-and-neck (1.1%) and esophageal (1.1%) cancers (Fig. 4C). Complete list of distribution per organ for FA pathway genes, *FANCA*, *FANCC*, *FANCD2, FANCE, FANCL*, *FANCF, FANCG, FANCI, FANCM and FANCL* are illustrated in Suppl. Fig. 10, Suppl. Table 1. Since in the clinical scenario genomic alterations in FA pathway have been most commonly considered for clinical trial inclusion in prostate cancer, we sought to analyze more in detail the prostate cancer cohort (n= 323). When looking at most common DNA repair genes within the prostate cancer cohort, concomitant alterations were identified most commonly in *BRCA2* (4.4%), *ATM* (5.0%), *CDK12* (4.3%) and *CHEK2* (1.2%) (Suppl. Fig. 9B). No differences regarding prevalence of concomitant DNA repair alterations were observed as compared to the FA pathway non-altered cohort. Together, our results establish *FANCA* as the most frequently altered FA pathway gene across human cancers and indicate that FA-defective tumors are characterized by increased genomic instability, while exhibiting heterogeneous DNA repair features. This underscores the need for alternative therapeutic strategies in this molecular subset.

**Figure 4.**
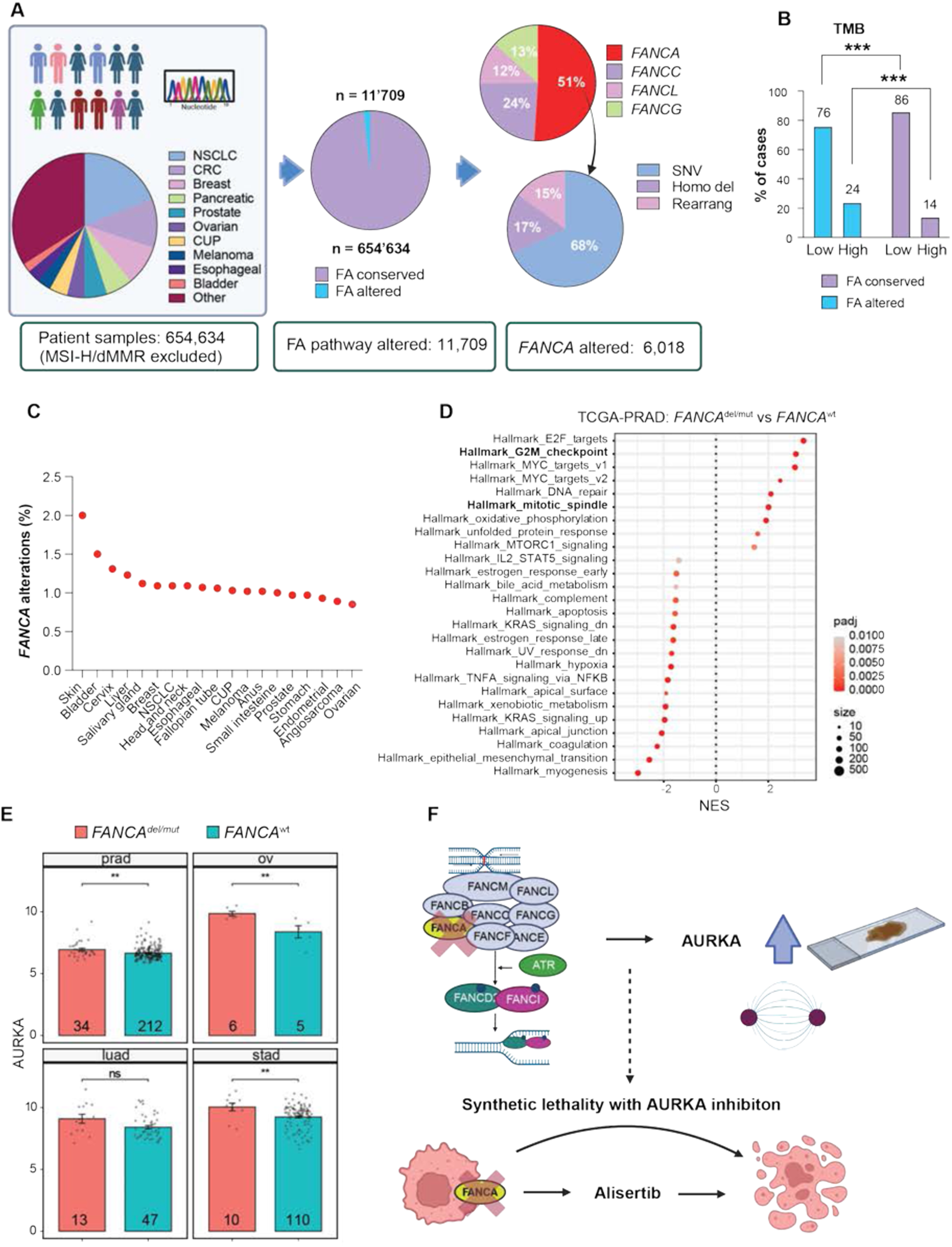
Genomic landscape of Fanconi anemia pathway gene alterations in human cancer. (A) Schematic illustrating Foundation Medicine NGS pan-cancer dataset analysis (FoundationOne^®^, FoundationOne^®^CDx and laboratory developed test FoundationOne^®^Heme). After excluding MSI-high/dMMR tumors, 654,634 patient tumor samples were available for analysis. Pathogenic or likely pathogenic FA pathway gene alterations were identified in 11,709 samples, of which 51% (n = 6,018) had alterations in *FANCA*. Relative frequency (%) of *FANCA*, *FANCC*, *FANCG*, and *FANCL* alterations and types of genomic alterations for the *FANCA* gene is illustrated. (B) Frequency of tumor with low vs high TMB among samples with and without FA pathway gene alterations: a higher proportion of FA pathway-altered tumors exhibit high TMB (24% vs 14%, p<0.001). (C) Percentage of tumor samples per organ harboring pathogenic or likely pathogenic genomic alterations in *FANCA*. (D) In silico analysis of RNA sequencing data from the prostate adenocarcinoma TCGA cohort confirms increased expression of the G2/M and mitotic spindle hallmarks in prostate adenocarcinoma samples with pathogenic or likely pathogenic *FANCA* genomic alterations, as compared to the FANCA- and *BRCA1/2*-wild type control cohort. (E) *AURKA* gene expression levels within the prostate cancer, ovarian cancer, lung adenocarcinoma, and stomach adenocarcinoma TCGA cohorts, comparison between *FANCA*-altered vs non-altered samples is illustrated. (F) FANCA deficiency is associated with AURKA upregulation/overexpression in human cancer, supporting the increased vulnerability of FANCA-deficient cancers to AURKA inhibition. *Abbreviations: CRC: colorectal cancer; CUP: cancer of unknown primary; FA: Fanconi anemia pathway alterations; NES: normalized enrichment score; NSCLC: non small-cell lung cancer; ov: ovarian cancer; padj: adjusted p value; prad: prostate adenocarcinoma, stad: stomach adenocarcinoma; wt: wild type.* Figure 4A *was created using BioRender*.

### AURKA expression in human cancer samples

To determine whether the AURKA dependency identified in functional screens is reflected in human tumors, we then examined AURKA expression and mitotic transcriptional programs in FANCA-altered cancer samples. Specifically, we performed an in silico analysis of the TCGA RNA sequencing dataset for five cancer types where pathogenic or likely pathogenic alterations in *FANCA* were most commonly identified: prostate, ovarian, breast, lung adenocarcinoma, and stomach cancers. Within the prostate and ovarian cancer datasets we observed an upregulation of the G2/M and mitotic spindle hallmarks in *FANCA*-altered vs non-altered samples (Fig 4D, Suppl. Fig. 9C). *AURKA* gene expression was upregulated within the prostate, ovarian and stomach cohorts (Fig 4E, Suppl. Fig. 9D). Within the ovarian, lung adenocarcinoma and prostate cohorts, we also observed upregulation of a series of centromeric and centrosomal genes (e.g., *CENPE, CENPF, KIF2C, KIF15, TTK*) (Suppl. Fig. 9D). When analyzing AURKA IHC levels in a cohort of FANCA-altered (n=5) vs non-altered (n=14) human cancer samples, we did not observe, however a characteristic differential pattern, possibly due to limited sample size and cancer type heterogeneity within the available tumor tissue cohort. However, we did observe a case with significant AURKA IHC staining (mean H-score 33) in a head-and-neck tumor sample with a pathogenic *FANCA* mutation p.R880* (c.2638C>T) (variant allele frequency 30%) (Suppl. Fig. 11A, B). Taken together, the analyzed patient data suggest that FANCA deficiency is associated with increased expression of AURKA and other mitosis-related genes in human cancer, supporting the findings from our screens.

## Methods

### Experimental models and cell culture

The DU145 (male, ATCC, HTB81™) and HEK293T cells (female, ATCC, RRID: CVCL_0063) cell lines were maintained in Dulbecco’s Modified Eagle Medium (DMEM) (Gibco) with 1% penicillin/streptomycin (P/S) and 10% fetal bovine serum (FBS). The PC3 (male, ATCC, RRID: CRL-3470) cells were cultured in RPMI 1640 medium (Gibco) with 1% P/S and 10% FBS. The cell lines CCH-SCC-FA1 (*FANCA*^⁻/⁻^),, CCH-SCC-FA1+Tg(LV) (*FANCA^Compl^*), UM_SCC_01 (*FANCA*^+/+^), and UM-SCC-01^KO^ (FANCA*^KO^*) were donated from the Fanconi Cancer Foundation and cultured in DMEM medium (Gibco), supplemented with 1% P/S, 10% FBS, and 1% non-essential amino acids (NEAA)^29^. RPE1 hTERT *p53*^-/-^, a kind gift from the Rottenberg lab, were cultured in DMEM medium (Gibco), supplemented with 1% P/S, 10% FBS, and 1% non-essential amino acids (NEAA). The cells were passaged twice per week using Phosphate Buffered Saline (PBS) and enzymatic detachment with TrypLE^TM^ Express Enzyme (Gibco). The medium was changed at regular intervals to maintain optimal nutrient supply, and the cells were observed microscopically to check for morphological changes and potential contamination. Cell culture was performed under standard conditions (37 °C, 5% CO2). Testing for mycoplasma contamination was performed regularly, at least twice per year.

#### Gene editing and silencing

##### Lentiviral transduction

Lentiviruses were produced by transient transfection of HEK293T cells with packaging plasmids and the corresponding plasmids using the lipofectamine 3000 transfection reagent (Thermo Fisher Scientific). After 30 h, lentivirus was collected from the supernatants on 2-3 consecutive days, concentrated 1:10 using the Lenti-X Concentrator (Takara), and resuspended in DMEM or PBS. Lentiviral stocks were stored at -80°. For lentiviral transduction, 250 μL of 10× concentrated lentivirus was added to a total volume of 5 mL medium on adherent target cells at a confluency of 60-70%, in the presence of Polybrene (Sigma) at a final concentration of 8 μg/mL. A second transduction was performed 24 hours later, and after 48 hours, cells were passaged. Antibiotic selection with blasticidin at a concentration of 10 μg/ml (Sigma), puromycin at 2-3 (20) μg/ml (Sigma), G418 (geneticin) at 500-1000 μg/ml (Thermo Fisher Scientific) and hygromycin at 500 μg/ml (Invivogen), was initiated 24 hours later an performed for at least 72 hours.

##### CRISPR/Cas9 knockout

For the knockout (KO) of *FANCA,* the DU145 and PC3 cell lines were transduced separately with 3rd-generation lentiviruses containing lentiCas9-Blast (Addgene Plasmid #52962) and gRNA targeting *FANCA* (Addgene Plasmid #111098). The cells were then selected using blasticidin at 10 μg/ml and puromycin at 2-3 µg/ml. After antibiotic selection, single-cell clones were derived by seeding single cells into at least 4 x 96-well plates using fluorescence-activated cell sorting (FACS). Single-cell-derived clones were expanded and screened for *FANCA* knockout by Western blotting.

##### Short hairpin (sh) RNA knockdown

*FANCA* shRNA knockdown (KD) was performed in RPE1 hTERT *p53*^-/-^ cells using two *FANCA*-targeting shRNAs (TRCN0000296799 and TRCN0000291182, Merck). Additionally, one non-targeting control shRNA (MISSION® TRC2 pLKO.5-puro, Merck) was used. Lentivirus generation, transduction, antibiotic selection, and WB to confirm *FANCA* KD was performed as described above.

##### Whole genome CRISPR/Cas9 synthetic lethality screen

On day 0, 8 million HEK293FT cells were seeded in 150cm cell culture dishes and on the following day transfected with lentiviral packaging plasmids and the Brunello all-in-one Hygromycin whole-genome library (CP1578, which targets 19,114 genes with a total of 76,7441 gRNAs), acquired from the Genetic Perturbation Platform (GPP, Board Institute), using 2xHBS (280nM NaCl, 100mM HEPES, 1.5mM Na2HPO4, pH 7.22), 2.5M CaCl2 and 0.1x TE buffer (10mM Tris pH8.0, 1mM EDTA pH8.0, diluted 1:10 with dH2O). After 30 h, virus-containing supernatant was collected during consecutive 5 days and concentrated using the LentiX concentrator (Takara), 1:10, and resuspended in PBS. For the multiplicity of infection (MOI) calculation, 150,000 target cells were seeded into 6-well plates. 24 hours later, virus at ascending MOIs was applied with 8 µg/ml Polybrene (Sigma), and 48 hours later, antibiotic selection with hygromycin at 500 µg/ml (Invivogen) was initiated and maintained for 7 days. For lentiviral transduction, 70 million cells per cell line were seeded in 15 cm dishes (5 million per dish) on day 0 of the screen, along with a control plate (one per cell line). On day 1, cells were transduced with concentrated lentivirus, resulting in a transduction efficiency of ∼20%. On day 2, the medium was changed to hygromycin (500 μg/ml; Invivogen). Hygromycin selection was performed for 7 days; afterwards, cells were reseeded for expansion for an additional 7 days. For the screen, 39 million cells per condition and replicate (2 replicates) were seeded in 15 cm dishes (3 million/15 cm dish) on day 0, for a coverage of 500 cells/sgRNA, given the negative selection screen design. Cells were collected and pelleted for DNA isolation on day 6. Genomic DNA was extracted with the MN Nucleo Bond XL Blood gDNA isolation kit (Macherey-Nagel, 740950.50). The entire screen was carried out in biological duplicate. PCR pre-check was performed before submitting for NGS to the Broad Institute. Sequencing of PCR-amplified sgRNAs was performed on a Novaseq SP (Illumina), followed by pooled sequencing data analysis with deconvolution of individual barcodes using the PoolQ software. Quality control of the screen was performed using R software. Results from two independent replicates were then analyzed using MAGeCK Robust Rank Aggregation (RRA) ^30^ and Maximum Likelihood Estimation (MLE) ^31^ modules. Replicate correlation was consistent, with R^2^ values above 0.92 in all samples. Correlation of changes between Day0 and Day7 was also consistent, with an R^2^ of 0.403 in *FANCA*^wt^ samples and 0.483 in *FANCA*^KO^ samples (Supp Fig. 1C). Hit identification was performed by comparing each Day7 to each corresponding Day0. Minimum effect size was delimited using either LFC (MAGeCK RRA) or Beta Score (MAGeCK MLE) of each distribution, as well as their difference to exclude correlating genes. For this, cutoffs were established at the distribution mean ± 2*SD. To identify *FANCA*^KO^ synthetic lethality targets, we focused on those genes that were in-between those cutoffs (*i.e.* not altered) in the *FANCA*^WT^ cells, but below the lower cutoff in the *FANCA*^KO^ cells. This allowed us to exclude essential genes and identify differential depletion of genes occurring only in the *FANCA*^KO^ cells, which points towards synthetic lethality. Downstream analysis consisted of weighted pathway enrichment analysis using hypergeometric testing with MAGeCKFlute^32^, using *FANCA*^KO^ MAGeCK RRA LFC from Day0 as the weights.

##### Semi-automated high-throughput drug screen

The semi-automated high-throughput drug screen was conducted in collaboration with NEXUS Personalized Health Technologies (ETH Zurich), a specialized high-throughput drug screening facility. DU145 *FANCA*^WT^ and isogenic *FANCA*^KO^ cells were screened with the MedChem Express Cell cycle/DNA damage library (Catalog HY-L004, 1624 compounds), at a concentration of 1µM. Drug testing was performed in a 384-well plate format using an integrated automation platform (HighRes Biosolutions). Briefly, cells were dispensed using a BioTek EL406 washer-dispenser and incubated for 24h before drugs were added. After a drug exposure time of 72 hours, CellTiter-Glo 2.0 was added, and cell viability was measured as luminescence readout (Tecan M1000 pro plate reader). Systematic variation from plate to plate was removed by standard normalization procedures. Assay quality was assessed based on the Z’-factor. Potential within-plate row-wise or column-wise stripe patterns and edge effects, which can arise during plate preparation, were eliminated using the median polish method of Tukey et al ^33^, or by subtracting a smooth polynomial using the loess function (Boutros). Differential activity was determined according to the workflow outlined by Prummer et al. ^34^.

#### Immunofluorescence

##### Micronuclei and multinuclei formation assays

Cells were cultured on coverslips (Epredia and Thomas Scientific) in 24-well plates, and exposed to alisertib (Selleck Chemicals) at a concentration of 500 nM for 48 hours, afterwards washed in PBS and fixed with 4% paraformaldehyde (PFA) for 20 min. Following fixation, cells were stained with antibodies diluted in staining buffer (SB), prepared with PBS, BSA 2% (Sigma), glycine 0.15% (Sigma) and Triton X-100 0.1% (Sigma). For CENP-B detection, cells were incubated with the primary rabbit anti-CENP-B antibody (dilution 1:50 in SB, ab259855, Abcam) for 1 hour at room temperature (RT), followed by washing and incubation with the Alexa Fluor™ 568 goat anti-rabbit antibody (dilution 1:1000 in SB, A11011, Invitrogen) for an additional 1 hour at RT. For Lamin-B1 detection, cells were incubated with the primary mouse anti-Lamin B1 antibody (dilution 1:200 in SB, MAB8525, Bio-Techne) for 1h at RT, followed by washing and incubation with the Alexa Fluor™ 488 goat anti-mouse antibody (dilution 1:1000 in SB, A11029, Invitrogen) for an additional 1 hour at RT. For α-tubulin detection, cells were incubated with the primary rat anti-α-tubulin antibody (Invitrogen, MA5-44046, 1:1000 in SB) for 1 hour at RT, followed by washing and incubation with the Alexa Fluor™ 568 anti-rat antibody (Invitrogen, A21247, 1:2000 in SB) for an additional 1 hour at RT. Finally, DAPI counterstaining was performed (10 mg/ml; Sigma, diluted 1:10’000 in SB) for 1 minute at RT, then washed with PBS. Images were acquired using a Nikon Ti2 spinning disk CREST Cicero. Z-stacks with at least 11 slices were performed at 10-30 spots per sample with a 60x oil objective. The slice thickness was approximately 3 μm. Micronuclei were counted semi-automatically using the Fiji ImageJ (2.0.0) plugin. Micronuclei and multinuclei were counted manually using the “Cell Counter” plugin in Fiji (ImageJ 2.0.0). Statistical significance was assessed using a one-way ANOVA in Prism 10 (10.6.1), followed by Šidák’s multiple comparison test.

##### FANCD2 foci formation assays

Cells were cultured on coverslips (Thomas Scientific) in 12-well plates and exposed to 1 μM Mitomycin C (Merck) for 24 hours; afterwards, they were washed in PBS and fixed with 4% PFA for 20 minutes on ice. Following fixation, cells were washed with PBS + 0.2% Tween 20 (Sigma) and then permeabilized for 20 minutes with PBS+ 0.2% Triton X-100 (Sigma) at RT. Afterwards, cells were washed twice with SB and blocked in the same SB for an additional 30 minutes at RT. Cells were then incubated with the primary antibody for 1 hour at RT, rabbit anti-FANCD2 antibody (Novus Biologicals, NB100-182, 1:500 in SB) or mouse anti-FANCD2 antibody (Novus Biologicals, NB100-316, 1:500 in SB). After 2 washes with SB, cells were incubated with goat anti-rabbit Alexa Fluor 647 (Jackson ImmunoResearch, 1:2500 in SB) for 1 hour at RT. After 5 additional washes with SB, cells were incubated with DAPI (10 mg/ml; Sigma, diluted 1:10’000 in SB) for 1 minute at RT, then washed again with PBS. Coverslips were fixed to microscopy slides (Meck Millipore) with Vectashield antifade mounting medium (AdipoGen). Images were acquired using a Zeiss LSM 710 (with Airyscan) or Zeiss LSM 980. Z-stacks with at least 5 slices were acquired at 5 spots per sample using a 63x oil objective; the stacks were then projected to a single layer for quantification. The slice thickness was around 3 μm. FANCD2 foci were then automatically counted using the Fiji image processing package in ImageJ (2.0.0). Briefly, cell nuclei were counted using the ‘‘analyze particles’’ order, and FANCD2 foci were identified and counted with the ‘‘find maxima’’ step. Statistical significance was assessed using an unpaired t-test in Prism 10 (10.6.1).

##### Cell cycle synchronization

For cell cycle synchronization in the G2/M phase, cells were treated with RO3306 (MedchemExpress) at 10 μM for 16 hours, then released into fresh, fully supplemented medium, with or without alisertib at 500nM. Drug treatment in synchronized cells was performed for 2 hours, overlapping with the release phase.

##### Immunohistochemistry (IHC) and IHC assessment

IHC stainings were performed by the Translation Research Unit (TRU) of the Institute of Tissue Medicine and Pathology at the University of Bern. Staining was performed on cell blocks or formalin-fixed paraffin-embedded (FFPE) tumor slides. Slides were stained with H&E and antibodies against FANCA and AURKA. The following antibodies have been used: anti-FANCA (Proteintech, catalog 11975-1-Ap; 1:1000) and anti-AURKA (35C1, Invitrogen, catalog 458900, 1:100 for FFPE material and 1:500 for cell blocks). IHC AURKA expression on FFPE tumor tissue sections was assessed by a board-certified pathologist (SdB) with Visiopharm software (v. 2025.02 x64, Horsholm, Denmark). Following procedures and workflow were used: tissue segmentation into tumor and non-tumor components, identification of regions of interest (ROI) using deep learning (DL)-based classification (U-Net); manual assessment and correction of ROIs where required; tumor cell segmentation into nuclear and cytoplasmic compartments using DL-based classification (U-Net), and classification of individual nuclei and cytoplasm as negative, weak, moderate or strongly AURKA positive based on the HDAB – DAB feature mean pixel values per object; 4) Generation of the following output: absolute counts for negative, weak, moderate vs strongly positive tumor cell nuclei and cytoplasm and calculation of H-scores (= (1 × % cells with weak staining) + (2 × % cells with moderate staining) + (3 × % cells with strong staining; range 0-300) for nuclei, cytoplasm and both combined. For all patient samples analysed by IHC, a signed general consent was available, and the study received approval from the local ethics committee (BASEC 2023-00976, Kantonale Ethikkommission Bern, Switzerland).

##### Western blotting

Cells were washed with PBS and lysed in RIPA buffer (Thermo Fisher Scientific) containing protease/phosphatase inhibitor cocktail (Thermo Fisher Scientific) for 20 minutes on ice. Afterwards, lysates were obtained by centrifugation (14’000 RPM, at 4°, 20 minutes). Protein concentration was measured using the Pierce BCA assay kit (Thermo Fisher Scientific), using a BSA standard curve. Protein lysates were denatured at 95°C for 5 minutes by adding 6x Laemmli SDS Sample Buffer (Thermo Fisher Scientific). Proteins were separated by Sure-page Bis-Tris gels 8% or 4-20% (GenScript) or Mini-Protean TGX Precast gels 4-15% (BioRad), then transferred by dry (iBlot2, Thermo Fisher Scientific, IB23001) transfer to nitrocellulose membranes (Sigma). Membranes were then blocked in 5% BSA (Sigma), dissolved in TBS-T (100 mM Tris, pH 7.5, 0.9% NaCl, 0.05% Tween-20), and incubated with primary antibodies dissolved in 5% BSA in TBS-T (4°C, overnight). After 3x washing in TBS-T, membranes were incubated in Horseradish Peroxidase (HRP)-linked (Cell Signaling, dilution 1:1000) or IRDye800CW (Licorbio, dilution 1:5000) secondary antibodies for 1 hour at RT. Western blot images were acquired using the Advansta enhanced chemiluminescent (ECL) detection on the FUSION FX7 EDGE Imaging System (Witec AG). For some experiments, the Odyssey Infrared Imaging System (LICOR) was used to acquire images. Quantification of western blots was performed using Fiji (ImageJ 2.16.0/1.54p). Quantification was performed in Fiji using the “Gels” analysis tool to calculate the area under the curve for each band, and normalized to the loading control. Following antibodies have been used: rabbit anti-AURKA (EP1008Y, ab52973, abcam, 1:1000), rabbit anti-vinculin (EPR8185, ab129002, abcam, 1:1000), rabbit anti-FANCD2 (EPR2302, ab221932, abcam, 1:1000), rabbit anti-FANCA (MA5-35800, Invitrogen, 1:1000).

##### Drug response assays

500 cells/well were seeded in 6 technical replicates on day 0; the next day, drugs were added at 6 or more concentrations, and cells were incubated with the drugs for 120 hours. Afterwards, cell viability was assessed with the Cell-Titer Glo 2.0 Reagent (Promega), and luminescence was measured on the Varioskan^TM^ LUX Multimode Microplate Reader (Thermo Scientific^TM^). Data were normalized to the corresponding untreated controls and analyzed with GraphPad Prism (version 10.5). We used a logarithmic or linear scale, depending on the concentrations range, to plot increasing drug concentrations on the X axis. Statistical significance was calculated using the 2-way ANOVA followed by Bonferroni’s multiple comparison test.

##### Clonogenic Growth Assays

Isogenic FANCA-proficient and -deficient cell lines (DU145_Cas9 and DU145_FANCA^KO^; CCH-SCC-FA1 (*FANCA*^⁻/⁻^) and CCH-SCC-FA1 (*FANCA^Compl^*) were seeded in 6-well adherent cell culture plates with 1000 cells/well for the DU145 cells and 2500-4000 cells/well for the CCH-SCC-FA1 cells, and exposed to target drugs, diluted to 3-4 distinct concentrations. Cells were seeded on day 0; drugs or DMSO were added on day +1; and cells were fixed after 12 days with 4% PFA (Sigma). Following fixation, colonies were stained with 0.1% crystal violet. Each experiment was performed with at least three biological replicates, each comprising two technical replicates. Controls were treated with DMSO. Scanned images were analyzed with ImageJ (Colony Area plugin) (2.0.0), and data were analyzed with GraphPad Prism (10.6.1), normalizing to the untreated controls. Statistical significance was calculated with 2-way ANOVA followed by Šidák’s multiple comparison test.

##### Flow cytometry and cell cycle analysis

CCH-SCC-FA1 (*FANCA*^⁻/⁻^) and CCH-SCC-FA1(*FANCA^Compl^*) cells were cultured in 10cm tissue-culture dishes (353003, Corning), and exposed to alisertib 500nM for 12, 24, 36, and 48h, or DMSO for 48h, afterwards washed in PBS. The cells were collected in 2µL Eppendorf tubes, washed once with PBS, and then fixed using 4% PFA for 20minutes on ice. Following fixation, cells were washed twice with TBS + 0.2% Tween 20 (Sigma) and then permeabilised for 20 minutes with TBS + 0.2% Triton X-100 (Sigma) at RT. Afterwards, cells were washed once with FACS buffer (TBS + 5% FBS). Cells were incubated with the primary rabbit anti-phospho-histone H3 (Ser10) (D2C8) monoclonal antibody (3377S, Cell Signaling Technology, 1:1200 in FACS Buffer) for 1 hour at RT. After washing twice with TBS-T, cells were incubated with anti-rabbit Alexa Fluor^TM^ 568 (A11011, Invitrogen, 1:1000 in FACS Buffer) and DAPI (D9542-10MG, Sigma-Aldrich, 1:10000 in FACS Buffer). After an additional wash with TBS-T, cells were resuspended in 1ml FACS buffer, transferred to round-bottomed tubes (352235, Corning), and sorted using the BD LSR II SORP flow cytometer (BD Biosciences). Generated data were processed with FlowJo software, v10.

##### Bulk RNA sequencing

CCH-SCC-FA1 (*FANCA*^⁻/⁻^) and CCH-SCC-FA1 (*FANCA^Compl^*) cells were treated for 12 and 48 hours with alisertib 500 nM or DMSO. Cells from 3 independent biological replicates were harvested, and total RNA extraction was performed using the Rneasy Mini Kit (Qiagen), including the recommended DNase treatment. RNA quantity and quality were analyzed using the De Novix DS-11 FX Spectrophotometer and Fluorometer (Labgene). RNA was then sent to Novogene for bulk sequencing. Messenger RNA was purified from total RNA using poly-T oligo-attached magnetic beads, and strand-specific and non-strand-specific libraries were created and pooled. Sequencing was performed in an Illumina NovaSeq X Plus at 30X sequencing depth. Quality control was performed with FastQC.

##### Bacterial culture, plasmid DNA isolation, and viral transduction

One Shot™ TOP10 Chemically Competent E.Coli (Thermo Fisher Scientific) were used for plasmid amplification. Briefly, 1 µl of plasmid was added to bacterial vials, which were heat-shocked at 42°C for 30 seconds, following the manufacturer’s instructions. Subsequently, 250 µl of S.O.C medium (Thermo Fisher Scientific) was added to the vials, which were then incubated in LB medium in a shaking incubator. Plasmid isolation was performed using the NucleoBond Xtra Midi EF (Macherey-Nagel), and plasmids were checked by electrophoresis and stored at -20°. Additionally, bacterial glycerol stocks were generated. For 3rd generation lentiviral production, HEK293T cells were grown in 10 cm dishes to 70-80% confluency in DMEM++. Before transfection, the medium was changed to Opti-MEM, and transfection was performed using Lipofectamine™ 3000 Transfection Reagent (Thermo Fisher Scientific). The following plasmid amounts were used: 7.5µg of plasmid of interest, 3.75µg of plasmid MDL,1.75µg of plasmid REV and 2 µg of plasmid VSVG.

##### RNA extraction and real-time quantitative reverse transcription PCR (RT-qPCR)

RNA isolation from cell pellets was performed using the Rneasy Mini Kit (Qiagen). Complementary DNAs (cDNAs) were generated using the FIREScript RT cDNA Synthesis Kit (Solis BioDyne), and the HOT FIREPol EvaGreen qPCR Mix (Solis BioDyne) was used to perform RT-PCR, following the manufacturer’s instructions. Quantitative RT-PCR assays were performed with three biological and three technical replicates, respectively, on the ViiA 7 system (Applied Biosystems) or the QuantStudio^TM^ 6 (Thermo Fisher Scientific). GAPDH was used as a housekeeping gene. The following primer sequences have been used: AURKA: GGA ATATGCACCACTTGGAACA (forward primer (FP)) and TAAGACAGGGCATTTGCCAAT (reverse primer (RP)); GADPH: GGT ATC GTG GAA GGA CTC ATG (FP) and ATG CCA GTG AGC TTC CCG TTC (RP).

##### Statistical analyses

Pooled sequencing data from the CRISPR screen were deconvoluted using the PoolQ tool (v. 3). Quality control was performed using R software, and analysis carried out using the MAGeCK Robust Rank Aggregation (MAGeCK RRA) and Maximum Likelihood Estimation (MAGeCK MLE) modules, downstream analysis using the R package *MAGeCKFlute*^30,35^. MAGeCK RRA and MLE modules were used to fit a negative binomial model using sgRNA abundance. Gene-level aggregation will be performed according to MAGeCK RRA/MLE. Calculations were performed on UBELIX ^36^, the HPC cluster at the University of Bern. Depleted sgRNAs were assessed as potential vulnerability hits. Overlap with targets identified within the drug screen was analyzed.

##### Bulk RNA sequencing analyses

Row reads were aligned against the human genome *hg38* reference using STAR v.2.7.11^37^. Gene quantification was performed using RSEM v.1.3.3 ^38^ with the “Aligned.toTranscriptome.out.bam” files from STAR as input. Gene annotations derived from GENCODE v48. RNA-seq data from TCGA were downloaded from the cBioPortal datahub ^39^. Cases were classified as *FANCA*-altered if they harboured pathogenic or likely pathogenic genomic alterations in *FANCA* (deep deletions, frameshift and premature stop mutations). Samples with missense mutations were excluded from the analysis, since the effect of these mutations on protein function remains unclear in many cases. Gene expression profiles of *FANCA*-altered tumors were compared with those of samples not carrying *FANCA, BRCA1 or BRCA2* alterations. Differential gene expression analysis was done in R v.4 using DESeq2 v.1.48.2 ^40^, and RSEM’s expected gene counts of coding genes as input (“data_mrna_seq_v2_rsem.txt” files for the TCGA datasets). Gene set enrichment analyses were done using the fgsea v.1.34.2 R package ^41^. Gene sets were downloaded from MsigDB ^42^.

##### NGS data analyses

We analyzed a total of 669,135 tumor samples that underwent targeted NGS profiling with one of the following Foundation Medicine (FMI) assays: FoundationOne^®^, FoundationOne^®^CDx or FoundationOne^®^Heme. Samples with microsatellite instability (MSI-high) were excluded for further analyses resulting in a total of 654,634 patient samples. Samples were profiled between December 2013 and September 2025. We interrogated the dataset for prevalence and concomitant genomic alterations for the following four FA pathway genes, *FANCA*, *FANCC*, *FANCG*, *FANCL*, which are covered by all the three above mentioned assays. Hybrid capture was performed to identify single nucleotide variants (SNV) and short insertions/deletions (short variants), copy number alterations (homozygous deletions and gene amplifications) and rearrangements. The TMB of ≥10 mutations per megabase (Muts/Mb) was classified as high (TMB-high). HRDsig is estimated from a machine learning based algorithm that predicts HRD status using a broad basket of copy number features, as described previously ^43^. Disease-specific co-occurrence and prevalence analyses were limited to tumor types with at least 100 total cases. Gene co-occurrence analysis was performed for genes altered in at least 10 samples, with additional FDR-based adjustment for multiple comparisons. Approval for this study, including a waiver of informed consent and the Health Insurance Portability and Accountability Act (HIPAA) waiver of authorization, was obtained from the Western Institutional Review Board (Protocol No. 20152817).

## Discussion

Uncovering successful molecularly tailored treatment strategies for FANCA-deficient cancers remains an unmet clinical need for patients and physicians. As we show in our analysis of a large pan-cancer NGS dataset, pathogenic or likely pathogenic genomic alterations in FA pathway genes are found across multiple cancer types, collectively affecting a substantial number of patients. Relevantly, more than half of these alterations occur in *FANCA*. However, when considering individual tumor entities, FANCA-altered tumors typically account for a small proportion of cases, often reported at <1% in most published cohorts. This results in limited representation in clinical studies and has precluded definitive conclusions regarding therapeutic benefit in this molecular subgroup. In the particular case of prostate cancer, patients with tumors harboring loss-of-function alterations in *FANCA* have been included in multiple PARP inhibitor trials (e.g., TOPARP-B, PROfound, MAGNITUDE, AMPLITUDE) ^1–3,5^. However, PARP inhibitors have repeatedly failed to demonstrate consistent benefit in this molecular patient population ^1–3,5^. Other therapeutic strategies evaluated *in vitro* to target FANCA-deficient cancers include ATM inhibitors. In this context, a genome-wide CRISPR screen performed in lung cancer cell lines showed that loss of FA genes was synthetically lethal with ATM inhibitors ^44^. Moreover, in FANCA-deficient cell lines, ATM inhibitors have shown preclinical activity ^44,45^. However, to our best knowledge, therapeutic targeting of ATM has not yet entered clinical trials for patients with *FANCA* loss-of-function alterations. Currently, in clinical practice, the finding of a *FANCA* genomic alteration on tumor NGS usually does not lead to a molecularly targeted treatment recommendation.

In the current study, we paired two orthogonal, highly complementary discovery platforms—a genome-wide CRISPR synthetic-lethality screen and a high-throughput small-molecule drug screen—to systematically identify actionable vulnerabilities in FANCA-deficient cancer cells. Intriguingly, both approaches independently converged on *AURKA* as a top candidate dependency. Pharmacological inhibition of AURKA led to pronounced CIN, as well as homogeneous and early cell cycle arrest in G2/M in FANCA-deficient cells, while FANCA-complemented cells largely remained in G1. These findings suggest that FANCA-deficient cells are less capable of buffering mitotic stress, resulting in heightened vulnerability to disruption of mitotic checkpoint control.

While the precise molecular mechanisms linking FANCA loss to AURKA dependency remain to be fully elucidated, our data support a model in which FANCA deficiency increases reliance on mitotic checkpoint integrity to preserve chromosome segregation fidelity. FANCA-deficient cells exhibit an elevated basal CIN and mitotic stress, accompanied by transcriptional upregulation of G2/M checkpoint and mitotic spindle gene programs, rendering them hypersensitive to pharmacological inhibition of AURKA activity. In this context, AURKA upregulation may reflect a compensatory response to chronic mitotic stress rather than a direct regulatory interaction with FANCA (Fig. 4F). Nevertheless, this state of heightened functional dependence on AURKA creates an exploitable therapeutic vulnerability.

Our findings further suggest that the vulnerability of FANCA-deficient cells is not limited to AURKA inhibition but extends to additional perturbations of mitosis, such as PLK1 inhibitors or IC261, which acts as a microtubule polymerization inhibitor ^26^. This is consistent with previous reports implicating FA pathway components in mitotic spindle regulation and chromosome segregation ^11,22,24^. Moreover, our *in vitro* validation experiments also confirmed that FANCA-deficient isogenic cells are more vulnerable to docetaxel, a microtubule-stabilizing chemotherapy drug ^46^. In our unbiased screening approach, we also observed an enrichment in genes controlling the G2/M transition other than *AURKA.* This included genes coding for checkpoint signaling members like ATR and proteins regulating mitotic DNA damage repair, such as members of the 9-1-1 complex and CIP2A. These proteins have recently gained interest for their role in safeguarding mitotic chromosomes from the consequences of DNA damage and replication stress ^47,48^. Satisfaction of the DNA damage checkpoint is one of the key steps for mitotic entry and is tightly regulated by AURKA and other mitotic kinases such as PLK1 ^25,49^. Together, these observations suggest that FANCA deficiency confers a broader mitotic dependency, rather than a vulnerability restricted to replication-associated DNA damage repair.

From a translational perspective, *FANCA* genomic alterations represent a readily identifiable biomarker through routine tumor sequencing, and AURKA expression or mitotic gene signatures may provide complementary stratification tools. To date, AURKA inhibitors, most frequently alisertib, have been assessed in clinical trials including a molecularly unselected patient population with advanced solid or hematological cancers, leading globally to only limited clinical efficacy ^50–52^. Currently, alisertib has an orphan drug designation from the Food and Drug Administration (FDA) for advanced and refractory small-cell lung cancer. Reported predictive biomarkers of response to alisertib include genomic alterations in *RB1*, *CDK6*, *RBL1/2*, as well as increased c-Myc tumor tissue expression ^53,54^. The phase 2 trial of alisertib in castration-resistant prostate cancer with neuroendocrine features observed a *FANCA* large deletion and AURKA overexpression in an exceptional responder to alisertib. The patient had an initial high tumor load with multiple lung and liver metastases, with confirmed small-cell histology, and remained 14 months in complete radiographic remission under treatment with alisertib ^50^. Two additional patients with maintained response in the trial had *AURKA* amplification and overexpression, respectively ^50^. Drug response modelling confirmed increased sensitivity to alisertib of the derived organoid ^50^. AURKA overexpression has been reported as independent adverse prognostic marker in many cancer types ^55–57^, and has been associated with resistance to PARP inhibitors ^58^. We hypothesize that the enhanced vulnerability we observed *in vitro* is at least in part due to this overexpression, which could be potentially exploited as an additional predictive biomarker. In addition, our observation that FANCA-deficient cells are hypervulnerable to docetaxel might be of particular clinical interest, since several clinical trials assessed the efficacy of AURKA inhibitors in combination with taxanes, either paclitaxel or docetaxel, however, in an unselected patient population ^54,59,60^. Moreover*, in vitro* studies in ovarian cancer suggested synergy with PARP inhibitors, combination that could be also further explored in FANCA-deficient cancers ^57^, in particular, since previous clinical trials in metastatic castration-resistant prostate cancers showed that responses to PARP inhibitors are quite heterogeneous and globally insufficient in FANCA-deficient cases ^1–3,5^.

Importantly, the therapeutic axis uncovered here differs from that targeted by PARP inhibitors. Unlike BRCA1/2-deficient tumors, our analysis shows that FANCA-deficient cancers do not uniformly display a profound HRD signature, which may explain the inconsistent benefit of PARP inhibition observed in clinical trials. Our data instead point to a vulnerability linked to mitotic progression and checkpoint control, highlighting a distinct biological dependency that is not captured by conventional DNA damage response–targeted strategies.

Our findings are supported by multiple orthogonal in vitro systems, including FA patient–derived cancer cell models, in combination with large-scale clinical genomics and transcriptomic analyses. Importantly, AURKA inhibitors such as alisertib have already undergone extensive clinical evaluation, lowering the barrier for rapid biomarker-driven translation in molecularly selected patient populations.

Taken together, our data support a model in which FANCA deficiency drives mitotic stress and checkpoint dependence, creating a synthetic lethal vulnerability to AURKA inhibition that is orthogonal to canonical DNA repair–targeted strategies. These findings provide a strong biological and translational rationale for biomarker-guided evaluation of AURKA inhibitors in FANCA-deficient cancers and extend the scope of precision oncology for this molecular subtype.

## Acknowledgements

We thank Agata Smogorzewska, M.D., PhD, Rockefeller University.

## Authors Contributions

D.A., M.G.F. and P.F designed the study and experiments; D.A., M.G.F., A.B., C.P., D.S.H.L., L.H., S.H., L.L., S.M., A.P.S., T.W., P.T., C.C. and S.D. performed experiments and analyses of results; D.A. and M.R. developed the concept, D.A. provided a clinical perspective to the results; D.A and E.V. provided material and/or clinical data; D.A., S.R, M.R., S.M and M.Ri provided administrative, technical and material support; D.A., M.G.F., A.B. and S.R. wrote the initial draft of the manuscript; M.F.G, A.B. and S.S performed the bioinformatics analyses; B.D. and S.d.B performed the pathology review and immunohistochemical evaluation; R.F., C.M-R and D.R. contributed to experimental design and analysis or results; C.Pi contributed to figure design, and all authors contributed to the final version of the manuscript.

## Supplementary figures

**Suppl Figure 1.**
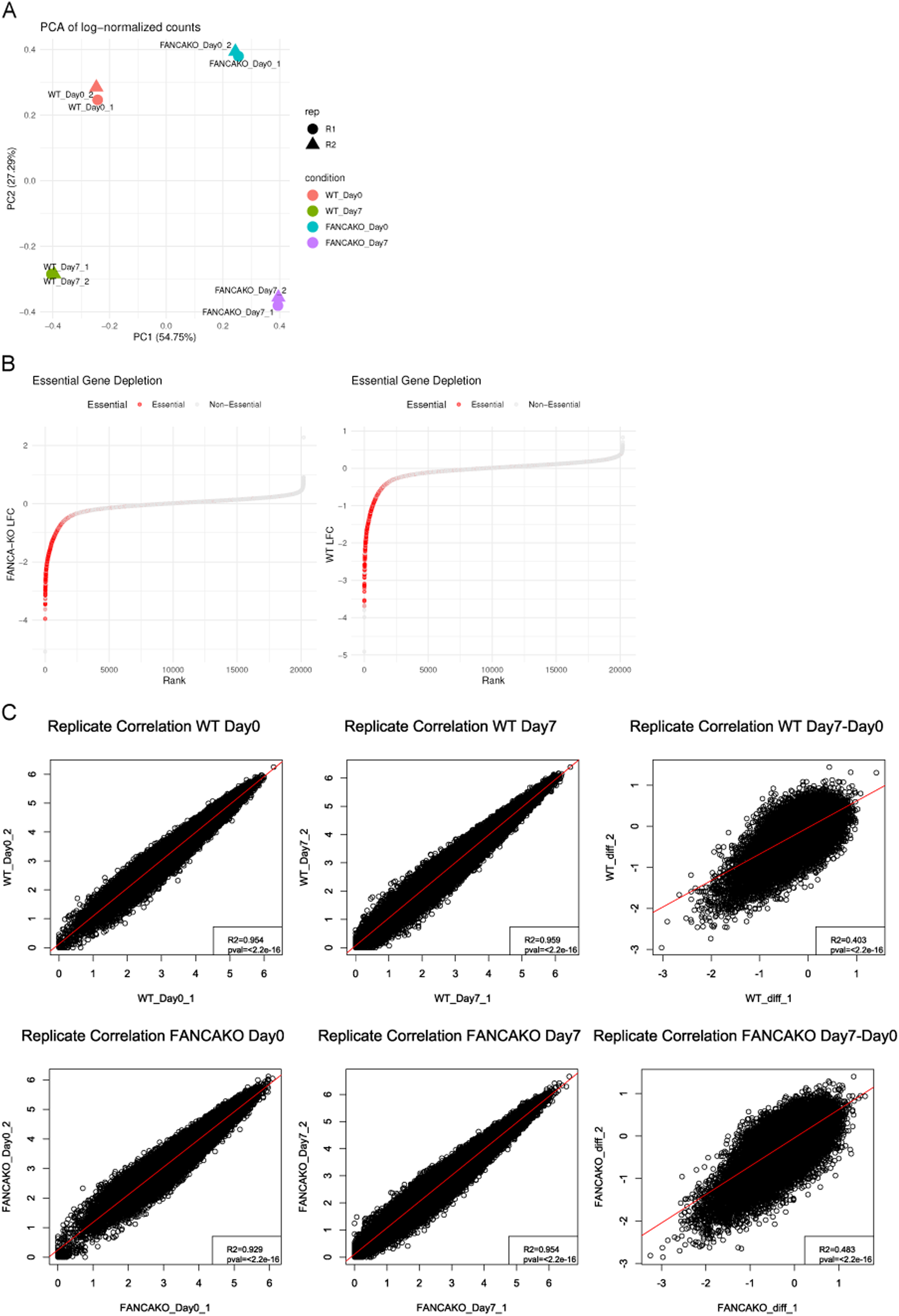
Quality control of the CRISPR/Cas9 synthetic lethality screen. (A) Principal Components Analysis (PCA) plot showing sample grouping and replicate correlation. (B) Ranked plot of gene LFC values in MAGeCK RRA in both *FANCA*^WT^ and *FANCA*^KO^ cells, highlighting essential gene depletion as a screen quality indicator. List of essential genes was obtained from Depmap 23Q2 public (Broad Insititute, 2023). (C) sgRNA count correlation plots for all CRISPR/Cas9 synthetic lethality screen samples showing good individual timepoint replicate correlation as well as the consistency of sgRNA abundance changes during the screen.

**Suppl Figure 2.**
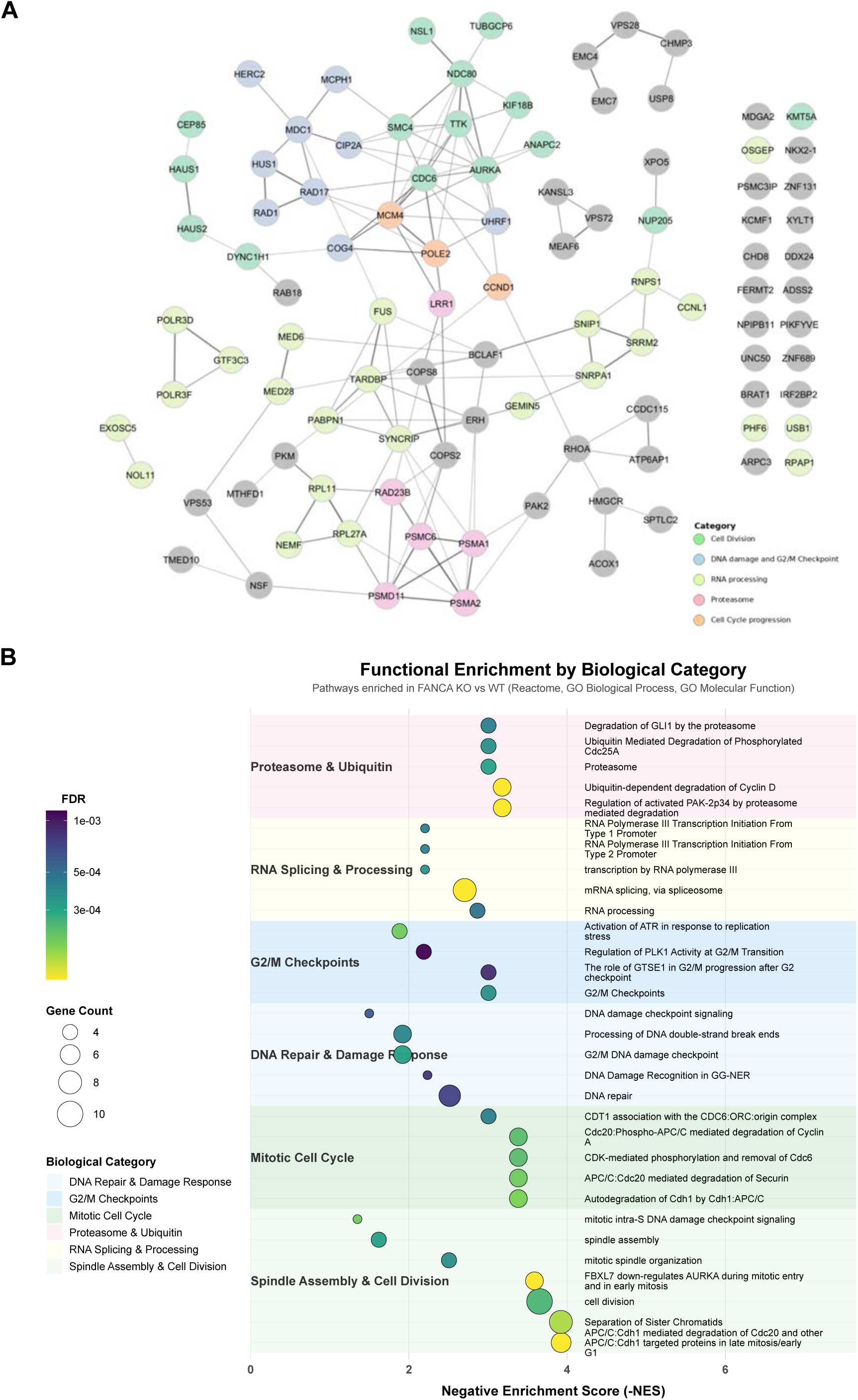
Pathway enrichment analysis of the hits identified in the CRISPR/Cas9 synthetic lethality screen. Whole genome loss-of-function screen was performed on isogenic FANCA -proficient vs -deficient DU145 cells. (A) STRING interaction map of the hits identified in the CRISPR/Cas9 synthetic lethality screen. (B) Extended pathway functional enrichment of the hits identified in the CRISPR/Cas9 synthetic lethality screen. Weighted hypergeometric analysis was performed on Gene Ontology and REACTOME databases.

**Suppl Figure 3.**
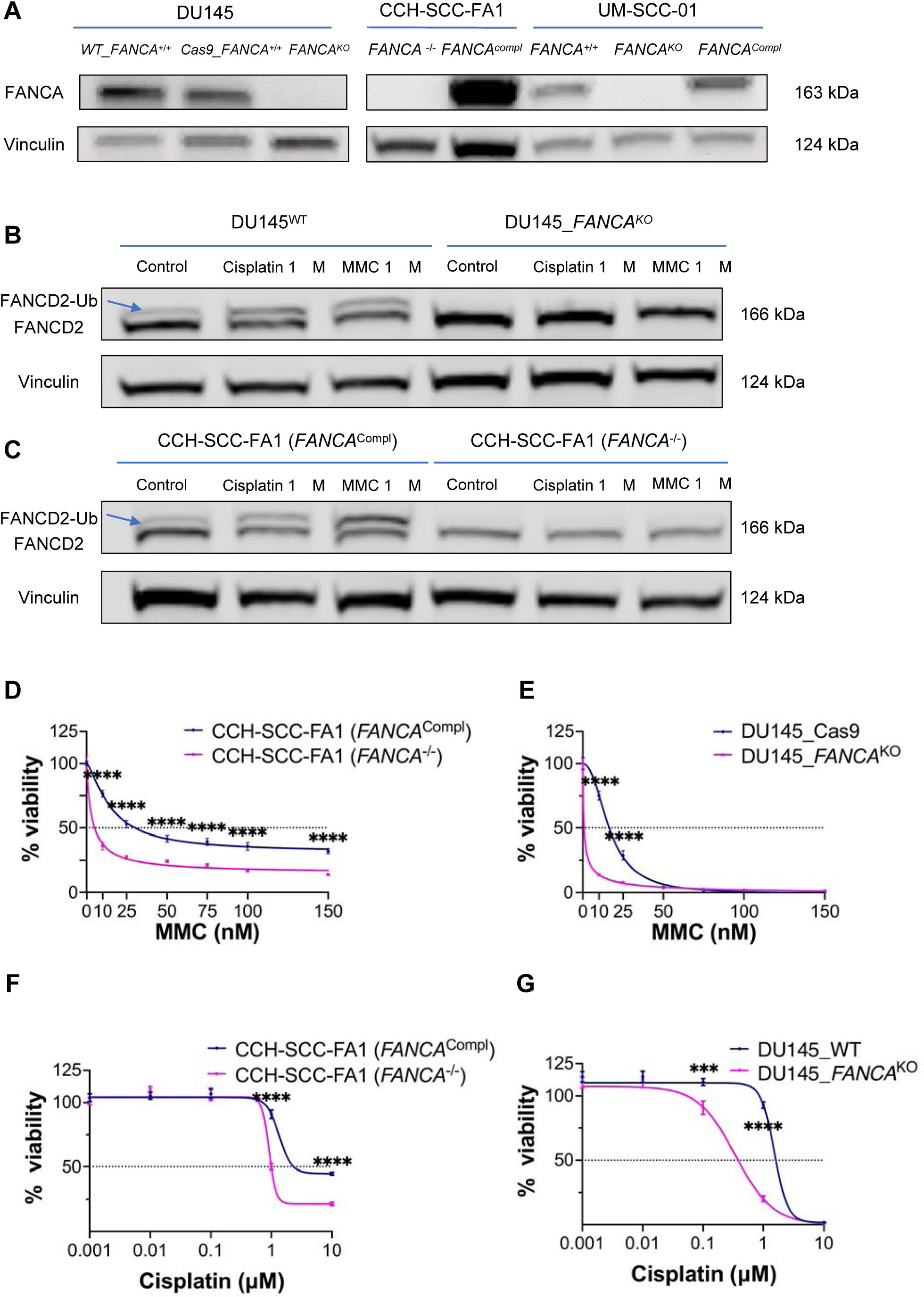
Extended data: functional characterization of FANCA-deficient vs proficient prostate adenocarcinoma and head-and-neck squamous carcinoma isogenic cell lines, including Fanconi anemia patient-derived cell lines. (A) Western blot illustrating FANCA protein loss in FANCA-deficient cell lines. (B,C) Lack of FANCD2 monoubiquitination, a hallmark of FA pathway activation, in FANCA-deficient cells. (D,E) FANCA-deficient cells show increased vulnerability to MMC: drug response curves for CCH-SCC-FA1 (*FANCA*^-/-^) vs CCH-SCC-FA1 (*FANCA*^Compl^) (D) and DU145^WT^ vs DU145_*FANCA*^KO^ (E) are illustrated. (F,G) FANCA-deficient cells show increased vulnerability to cisplatin: drug response curves for CCH-SCC-FA1 (*FANCA*^-/-^) vs CCH-SCC-FA1 (*FANCA*^Compl^) (F) and DU145^WT^ vs DU145_*FANCA*^KO^ (G) are illustrated.

**Suppl Figure 4.**
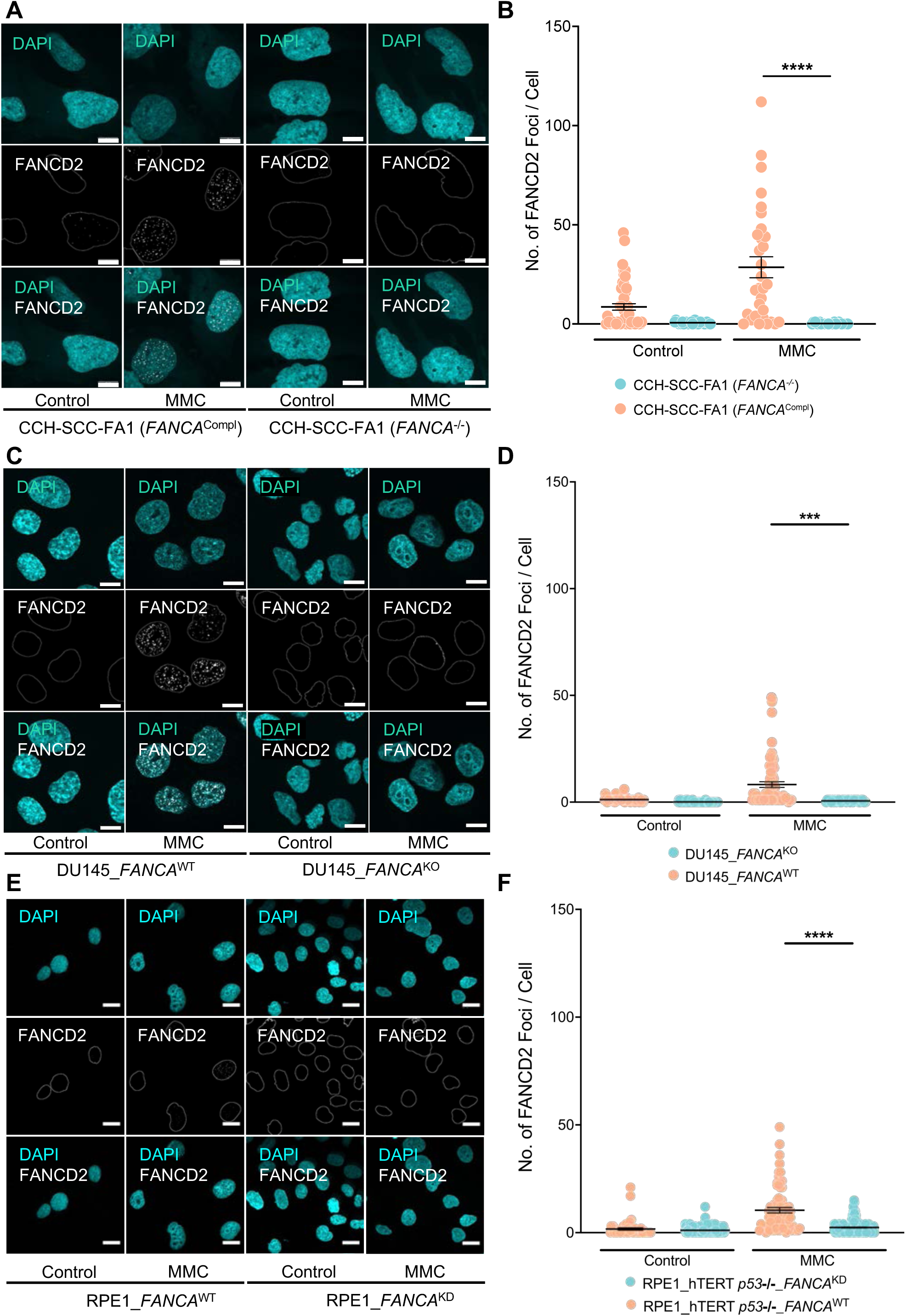
Extended data: FANCD2 foci formation in FANCA-deficient vs proficient isogenic cell lines under treatment with MMC and in control conditions. Cells were exposed to 1 µM MMC for 24 hours vs control conditions (DMSO) and imaged via confocal microscopy using Zeiss LSM 710 or Zeiss LSM 980 (63x). (A,C,E) Representative immunofluorescence images for CCH-SCC-FA1 (*FANCA*^Compl^) vs CCH-SCC-FA1 (*FANCA*^-/-^) (A), DU145^WT^ vs DU145_*FANCA*^KO^ (C), and RPE1^WT^ vs RPE1_*FANCA*^KD^ (E); scale represents 20 µm. (B, D, F) FANCD2 foci quantification for the same FANCA-deficient vs proficient cell lines. The number of foci per cell are represented, and statistical significance was calculated with the unpaired t-test.

**Suppl Figure 5.**
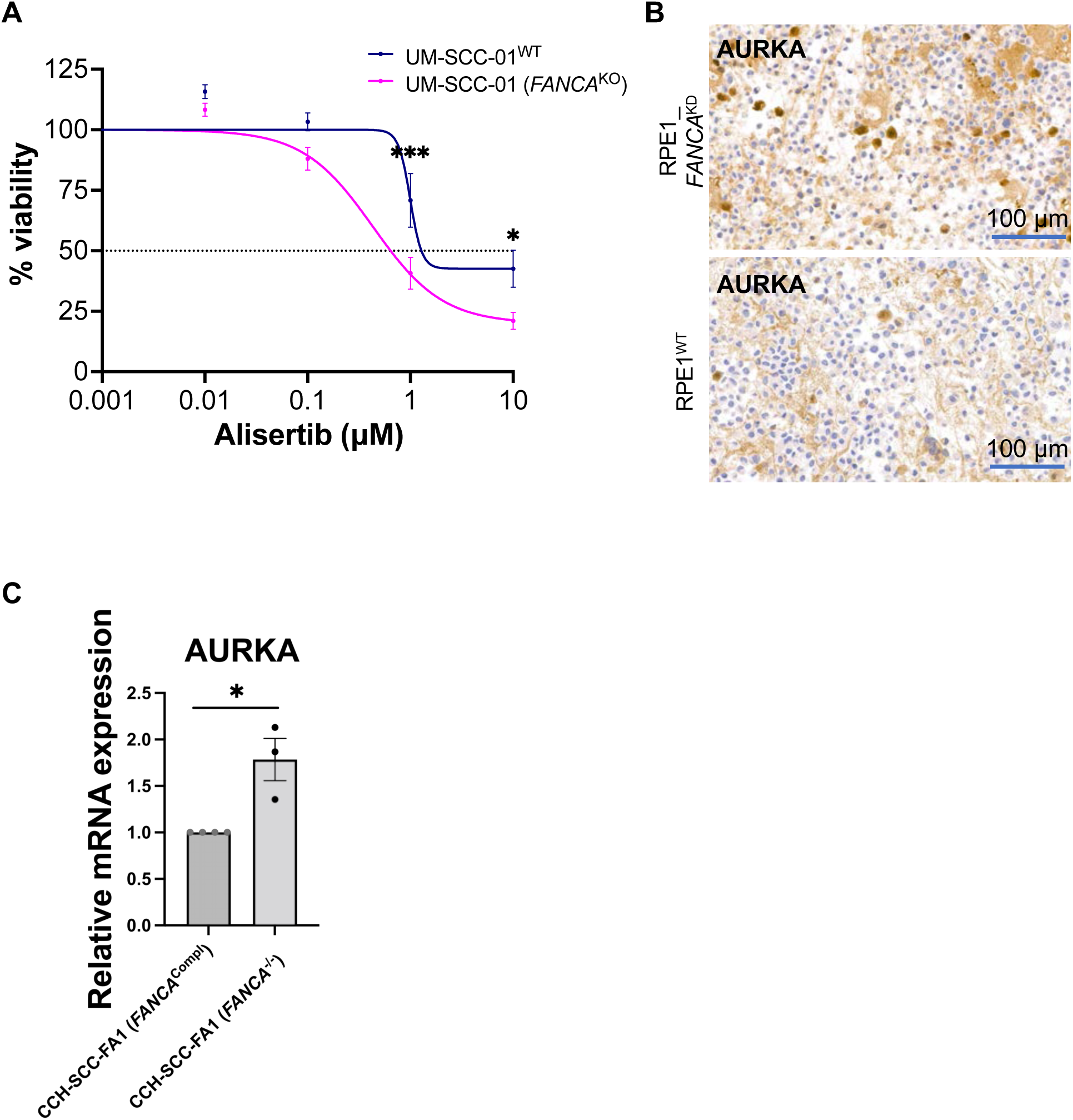
Extended data: *In vitro* hit validation from CRISPR/Cas9 whole-genome synthetic lethality and high-throughput drug screens confirms that FANCA loss is synthetically lethal with AURKA inhibition *in vitro*. (A) Drug response curves, assessed by CellTiter-Glo luminescence cell viability assay, illustrating increased vulnerability to the AURKA inhibitor alisertib of FANCA-deficient squamous head-and-neck cancer cell lines UM-SCC-01 (*FANCA*^WT^), as compared to the isogenic *FANCA*^KO^ cells, UM-SCC-01 (*FANCA*^-/-^). Statistical significance was calculated using the 2-way ANOVA followed by Bonferroni’s multiple comparison test. (B) Increased AURKA protein IHC expression shown on RPE1_*FANCA*^KD^ vs RPE1^WT^ cell lines, scale bar represents 100 µm. (C) Basal relative *AURKA* mRNA expression for the CCH-SCC-FA1 (*FANCA*^-/-^) vs CCH-SCC-FA1 (*FANCA*^Compl^) cell lines shows increased *AURKA* expression in FANCA-deficient cells. Statistical significance was calculated using the Mann-Whitney U test.

**Suppl Figure 6.**
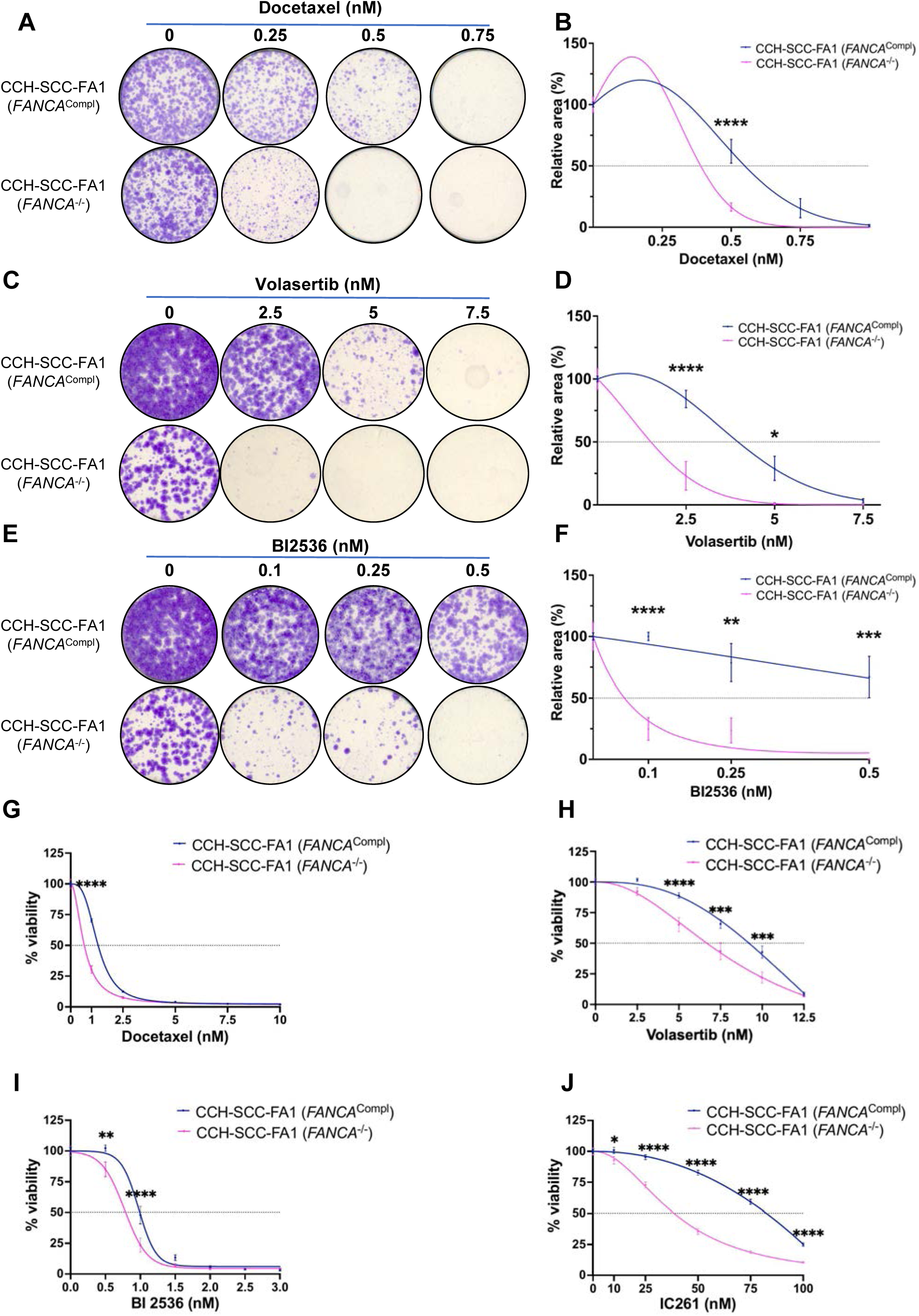
Extended data: *in vitro* hit validation from CRISPR/Cas9 whole-genome synthetic lethality and high-throughput drug screens for additional mitosis-regulating drugs. (A-F). Clonogenic growth assays for FANCA-deficient vs proficient isogenic CCH-SCC-FA1 (*FANCA*^-/-^) vs CCH-SCC-FA1 (*FANCA*^Compl^) cells under treatment with docetaxel (A,B), volasertib (C,D), and BI2536 (E,F): representative images (A,C,E) and relative area (B,D,F). Statistical significance was calculated with 2-way ANOVA followed by Šidák’s multiple comparison test. (G-I) Drug response curves illustrating increased vulnerability to docetaxel (G), volasertib (H), BI2536 (I) and IC261 (J) of FANCA-deficient vs proficient isogenic CCH-SCC-FA1 (*FANCA*^- /-^) vs CCH-SCC-FA1 (*FANCA*^Compl^). Statistical significance was calculated using the 2-way ANOVA followed by Bonferroni’s multiple comparison test.

**Suppl Figure 7.**
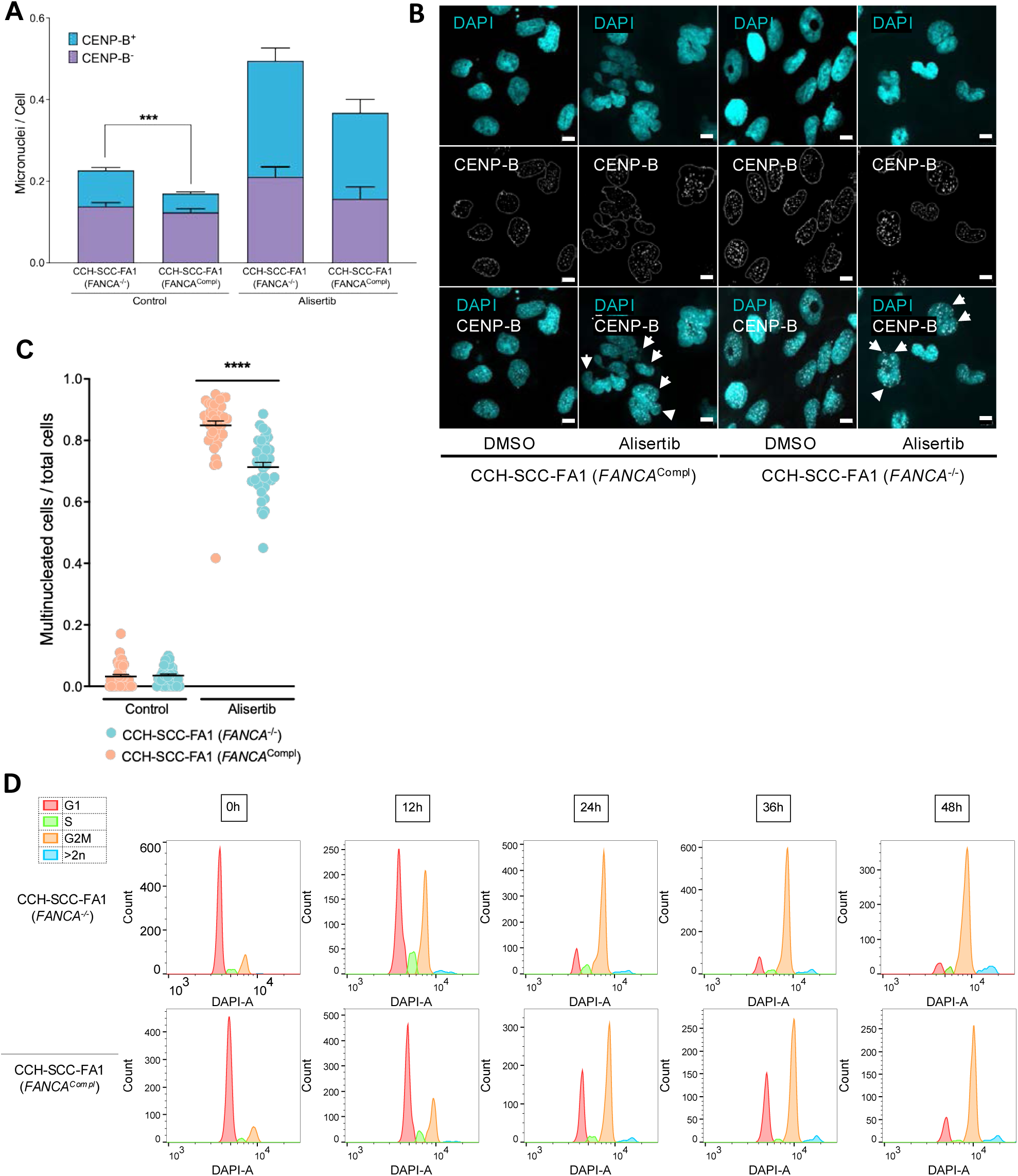
Extended data: in vitro validation of vulnerability to alisertib. (A) Quantification of CENP-B foci in micronuclei under treatment with alisertib (500 nM) vs. control conditions: the number of foci per non-multinucleated cells is represented; statistical significance refers to proportion of CENP-B positive micronuclei and was calculated with the one-way ANOVA test followed by Šidák’s multiple comparison test. (B-C) Quantification and (B) and representative images (C) of multinucleated cells under treatment with alisertib (500 nM) vs control conditions. Scale represents 10 µm. Statistical significance was calculated with the one-way ANOVA test followed by Šidák’s multiple comparison test. (D) Representative biological replicate from cell cycle analysis under treatment with alisertib 500 nM in CCH-SCC-FA1 (*FANCA*^-/-^) vs CCH-SCC-FA1 (*FANCA*^Compl^) cells.

**Suppl Figure 8.**
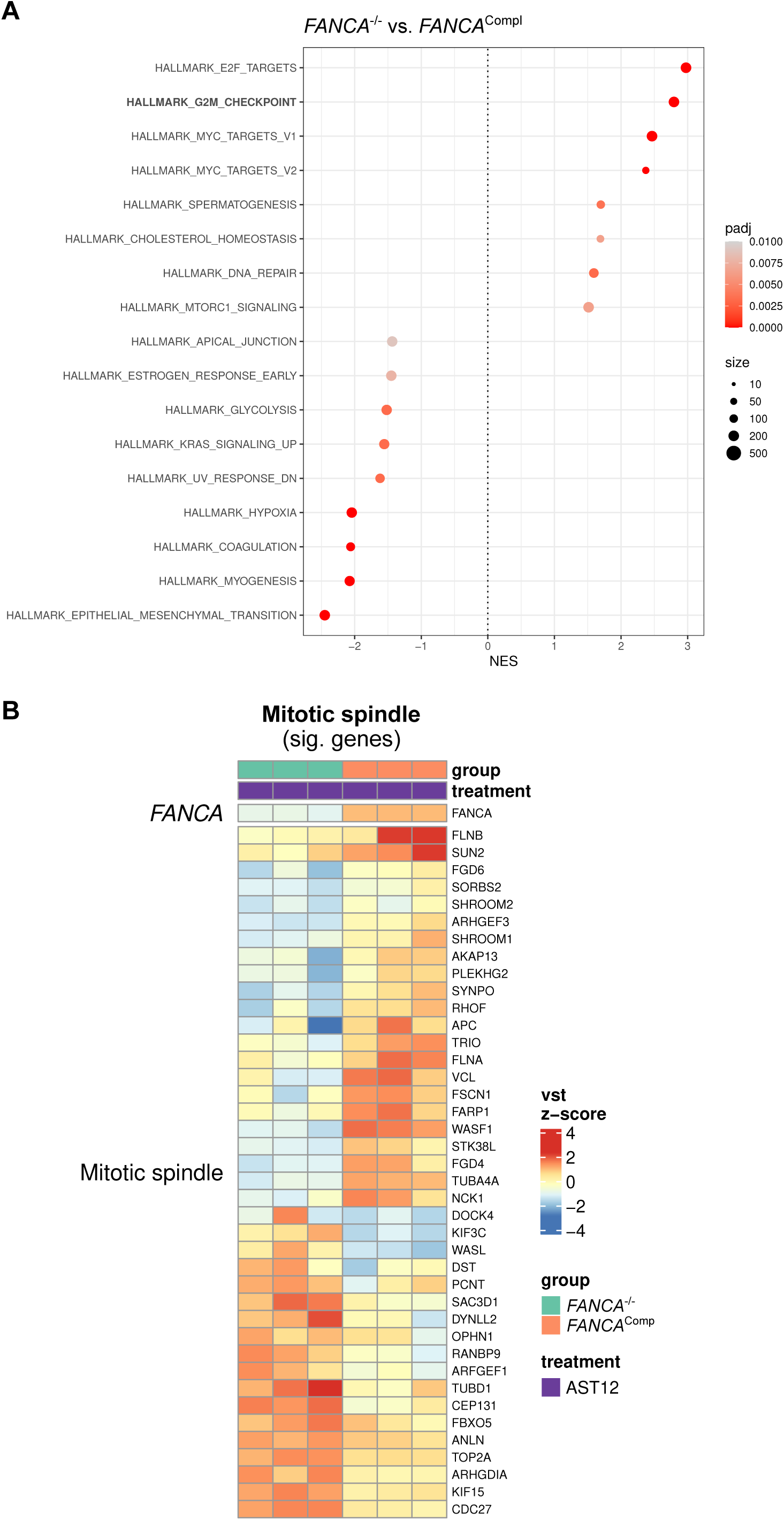
Extended data: in vitro validation of vulnerability to alisertib, part 2. (A) Functional enrichment analysis of bulk RNA sequencing from CCH-SCC-FA1 (*FANCA*^-/-^) vs CCH-SCC-FA1 (*FANCA*^Compl^) cells shows increased expression of the G2/M hallmark; basal levels are shown. (B) Heatmap illustrating individual genes from the mitotic spindle hallmark for CCH-SCC-FA1 (*FANCA*^-/-^) vs CCH_SCC_FA1 (*FANCA*^Compl^) cells after 12-h treatment with alisertib: upregulation of centrosome-involved genes is observed for the FANCA-deficient cells (*CEP 131, KIF15, TOP2A*). *Abbreviations: AST: alisertib; FA neg.: CCH-SCC-FA1 (FANCA^-/-^), FA pos.: CCH-SCC-FA1 (FANCA*^Compl^*), NES: normalized enrichment score; padj: adjusted p value.*

**Suppl Figure 9.**
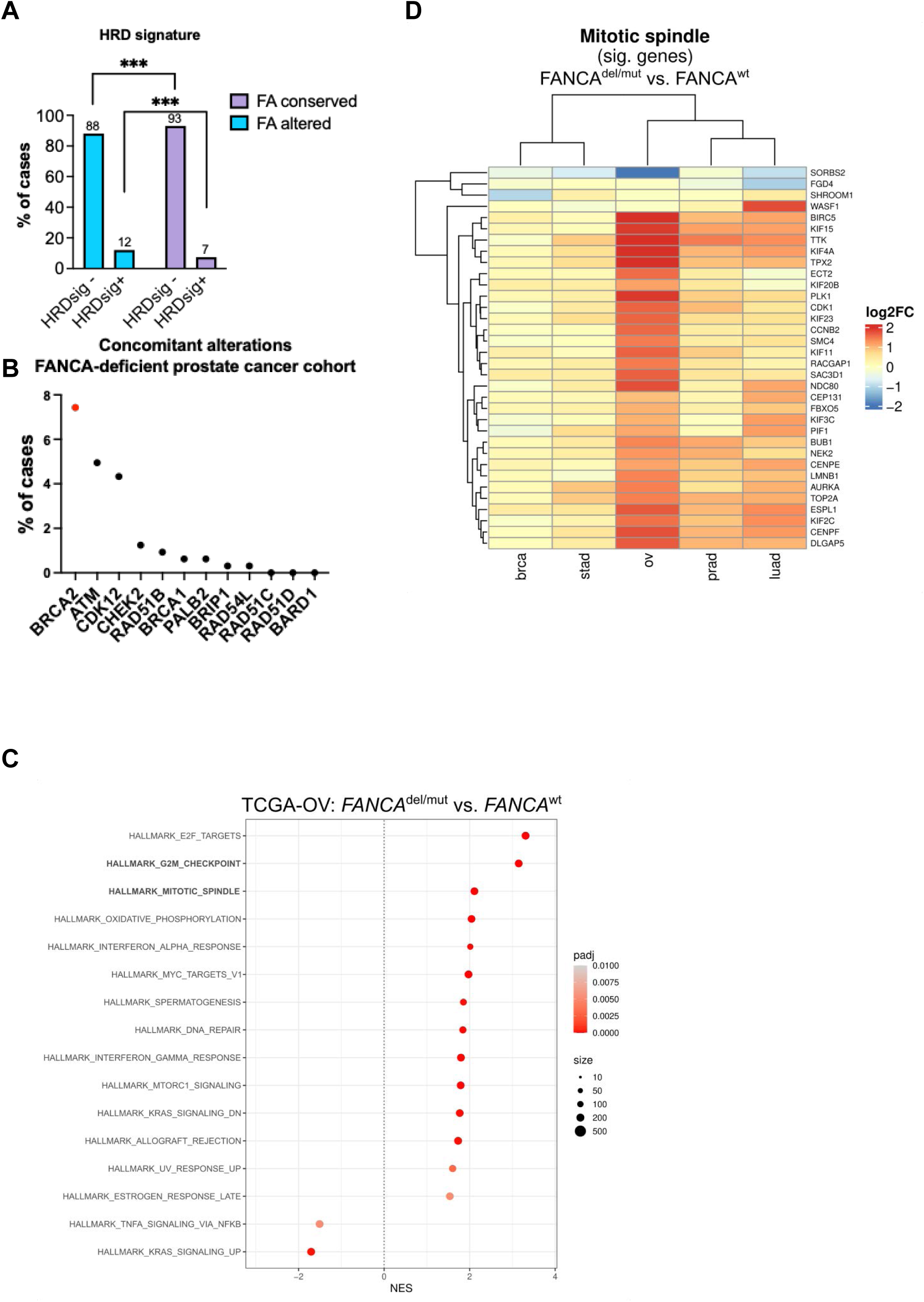
Extended data: genomic landscape of Fanconi anemia pathway gene alterations in human cancer. (A) Frequency of tumor with negative vs positive HRD signature among samples with and without FA pathway gene alterations: a higher proportion of FA pathway-altered tumors exhibit a positive HRD signature (12% vs 7%, p<0.001). (B) Concomitant alterations in other DNA repair genes within the FANCA-altered prostate cancer FMI cohort. (C) In silico RNA sequencing analysis from the TCGA ovarian cancer cohort shows increased expression of the G2/M and mitotic spindle hallmarks in ovarian cancer samples with pathogenic or likely pathogenic *FANCA* genomic alterations, as compared to the *FANCA* and *BRCA1/2*-wild type control cohort. (D) Heatmap illustrating individual gene expression within the mitotic spindle hallmark, including *AURKA*, in *FANCA*-altered vs wild-type TCGA tumor samples. Comparative gene expression profiles for prostate, ovarian, breast cancers, stomach, and lung adenocarcinoma are shown. *Abbreviations: FA: Fanconi anemia pathway; HRD: homologous recombination deficiency, NES: normalized enrichment score; ov: ovarian cancer; padj: adjusted p value; wt: wild type.*

**Suppl Figure 10.**
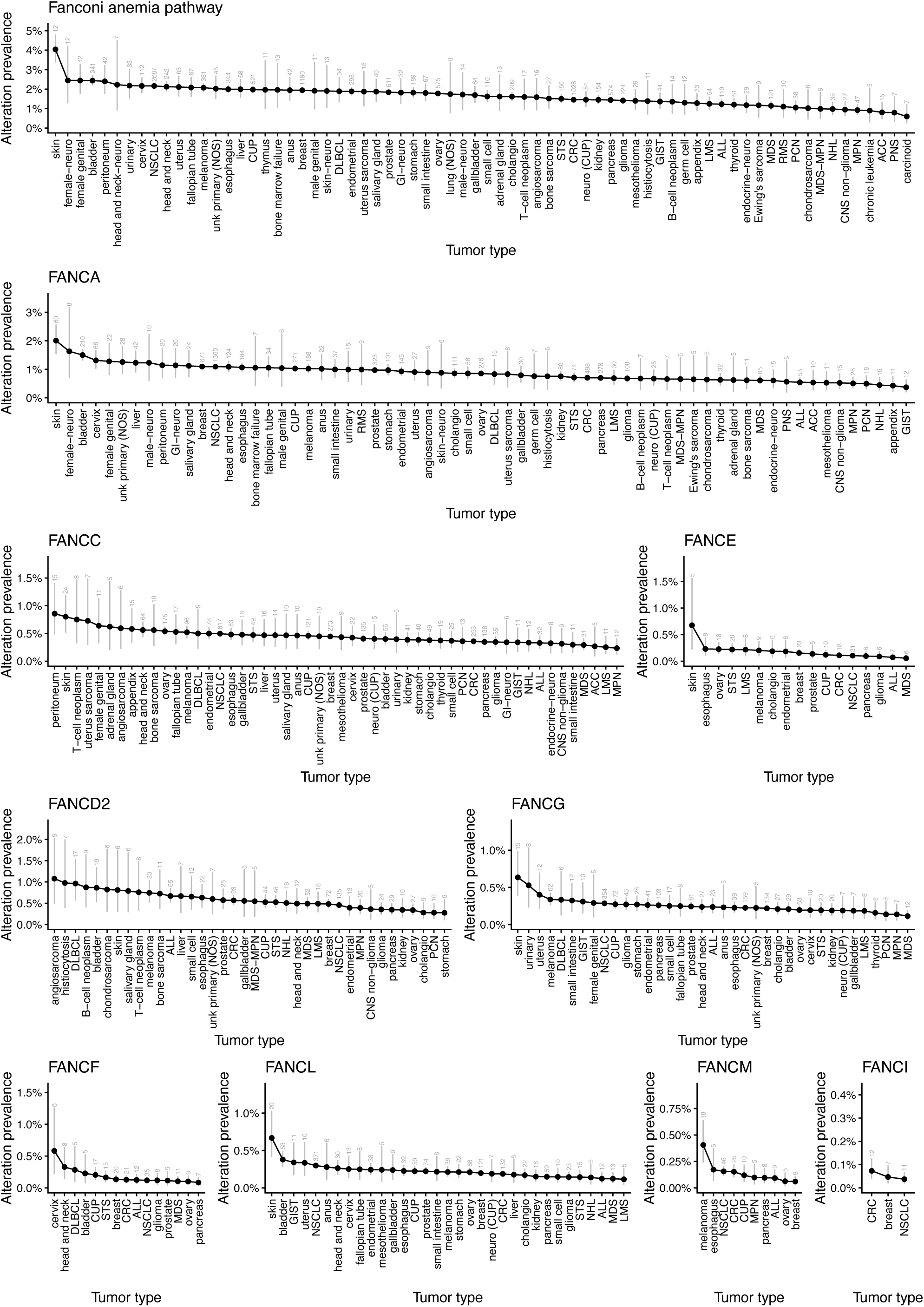
Frequencies by organ/tumor site for FA pathway gene alterations including distribution for individual gene alterations are illustrated.

**Suppl Figure 11.**
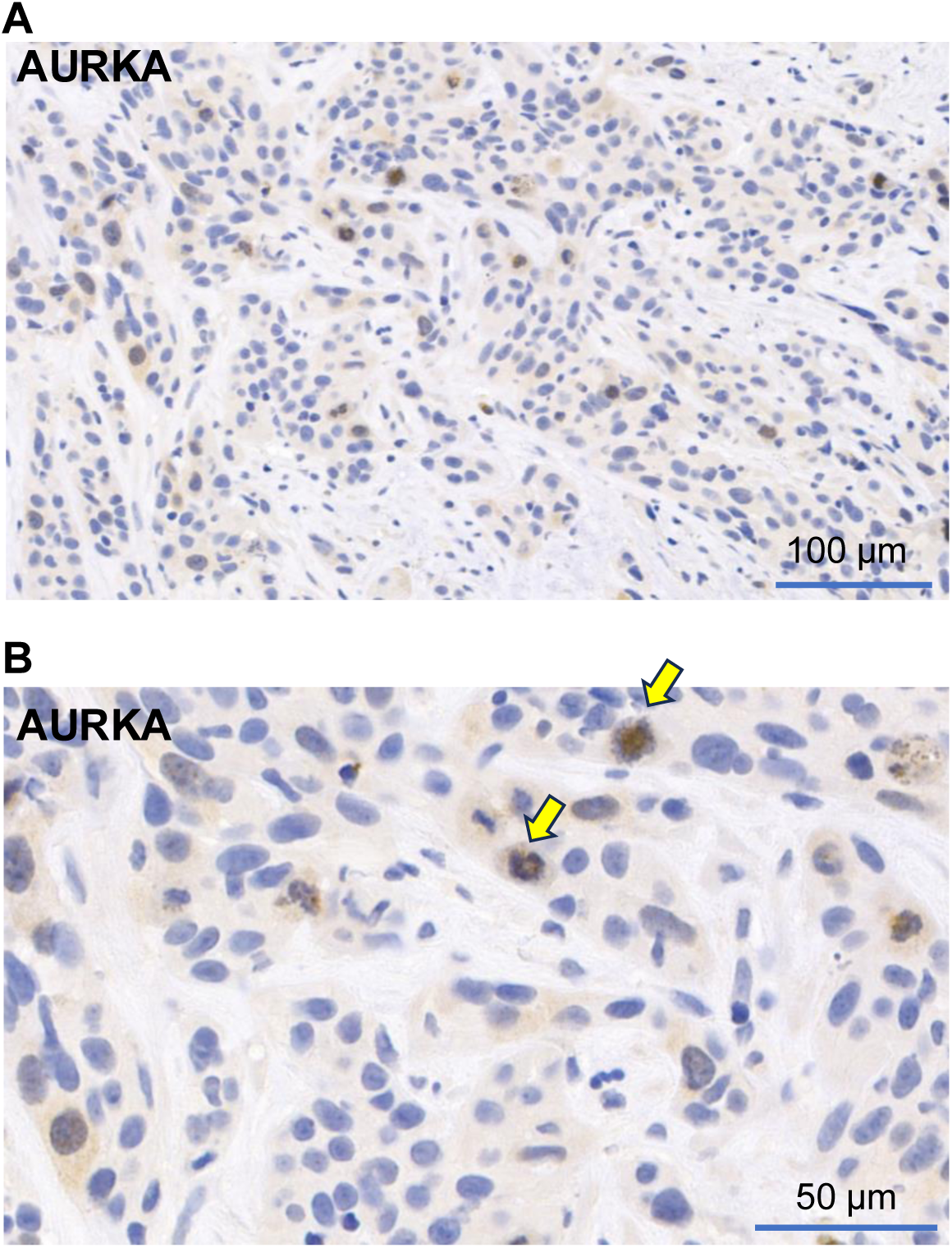
AURKA IHC staining on a head-and-neck squamous *FANCA*-mutated (p.R880* (c.2638C>T), VAF 30%) carcinoma FFPE sample. (A,B). Diffuse, weak cytoplasmic and occasional nuclear staining (mean H-score nuclear/cytoplasmic combined: 33), as well as specific moderate to strong centrosomal staining in mitotic cells (yellow arrows, B), is observed.

